# Ligandability at the membrane interface of GPx4 revealed through a reverse micelle fragment screening platform

**DOI:** 10.1101/2024.05.09.593437

**Authors:** Courtney L. Labrecque, Brian Fuglestad

## Abstract

While they account for a large portion of drug targets, membrane proteins (MPs) present a unique challenge for drug discovery. Peripheral membrane proteins (PMPs), a class of proteins that bind reversibly to membranes, are also difficult targets, particularly those that function only while bound to membranes. The protein-membrane interface in PMPs is often where functional interactions and catalysis occur, making it a logical target for inhibition. However, interfaces are underexplored spaces in inhibitor design and there is a need for enhanced methods for small-molecule ligand discovery. In an effort to better initiate drug discovery efforts for PMPs, this study presents a screening methodology using membrane-mimicking reverse micelles (mmRM) and NMR-based fragment screening to assess ligandability in the protein-membrane interface. The proof-of-principle target, glutathione peroxidase 4 (GPx4), is a lipid hydroperoxidase which is essential for the oxidative protection of membranes and thereby the prevention of ferroptosis. GPx4 inhibition is promising for therapy-resistant cancer therapy, but current inhibitors are generally covalent ligands with limited clinical utility. Presented here is the discovery of non-covalent small-molecule ligands for membrane-bound GPx4 revealed through the mmRM fragment screening methodology. The fragments were tested against GPx4 in bulk aqueous conditions and displayed little to no binding to the protein without embedment into the membrane. The 9 hits had varying affinities and partitioning coefficients and revealed properties of fragments that bind within the protein-membrane interface. Additionally, a secondary screen confirmed the potential to progress the fragments by enhancing the affinity from > 200 µM to ∼15 µM with the addition of certain hydrophobic groups. This study presents an advancement of screening capabilities for membrane associated proteins, reveals ligandability within the GPx4 protein-membrane interface, and may serve as a starting point for developing non-covalent inhibitors of GPx4.

## Introduction

Membrane proteins (MPs) account for around 23% of the proteome and over 60% of current drug targets.^1^ Peripheral membrane proteins (PMPs) are a diverse subcategory of water soluble, membrane associated proteins that interact, often reversibly, with membranes via electrostatic, hydrophobic, or specific lipid headgroup interactions.^2,3^ PMPs have emerged as a prominent category of drug targets for a variety of disease processes.^2,4^ Despite the great need for inhibitor development for integral membrane proteins and PMPs, many roadblocks remain.

While integral membrane proteins do present a challenge for certain inhibitor design approaches, high-throughput screening (HTS) and some structure-based design methods have had success, with the majority of small-molecule inhibitors confined to aqueous-exposed binding sites.^1,5^ Recently, attention has shifted towards inhibitors that bind within the protein-membrane interface, with this mode of inhibition becoming more prominant.^6^ Despite the recognition of the importance of drug-lipid interactions, protein-lipid interfaces are oftentimes overlooked during the initial stages of screening and rational drug design, due to challenges in housing proteins in a membrane-like environment that is also compatible with screening.^7–9^

Like integral membrane proteins, PMPs are also challenging drug targets. This class of protein is oftentimes only active when bound to the membrane, with the water-solubilized state being inactive.^4^ For many PMPs, the membrane interface generally represents the functional site of the protein that should be targeted for inhibition.^4^ Despite this, PMPs are typically screened in bulk aqueous conditions due to the readily available methods and the retained stability of this class of proteins in the absence of a membrane.^4,10^ To date, many PMPs are considered “undruggable,” meaning, the target is considered too difficult to probe by standard methods of screening.^1,11,12^ A hallmark of druggability is the presence of a solvent-accessible hydrophobic pocket, often but not exclusively, represented by an active site for enzyme targets.^1,13^ On the other hand, when the target is a membrane-associated protein, active sites do not necessarily translate to solvent-accessible.^1^ To access a specific binding site of a membrane protein that is exposed to the lipid phase, the small molecule will first have to partition into or onto the bilayer before engaging binding sites on the target.^6,7^ The ability of such sites to bind small molecules is termed ligandability, which is a prerequisite for a target to be considered druggable.^14,15^ Ligandability of membrane associated proteins at the membrane is an underexplored area, meaning it is mostly unknown the types of ligands that could target protein-membrane interfaces.^6^

A powerful approach for inhibitor discovery and design, fragment-based drug discovery (FBDD), shows great promise for discovery of new modes of inhibition. FBDD allows discovery of novel compound classes since the fragment libraries are typically optimized for chemical diversity and nearly infinite possibilities of combinations provide ample pathways to previously unknown inhibitor types.^19,20^ FBDD is based on initial discovery of very small molecules (< 300 Da) that bind to a protein target.^21^ Fragment screening is a useful approach for assessing whether a protein target is ligandable and reveals the initial building blocks for FBDD approaches.^14,22^ Due to the small size, two or more fragment hits may be linked, fragments may be grown, or elements of fragments can be combined to yield a larger, higher affinity compound.^23^ Due to the inherently weak nature of fragment binding, biophysical screening methods are almost always required.^23–25^ Additionally, optimal elaboration generally requires structural information, which can be provided by protein-detected NMR as well as crystallography.^26,27^ Widely used membrane models for biophysical methods include micelles, bicelles, liposomes, and nanodiscs, with the recently developed membrane-mimicking reverse micelles (mmRMs) as a promising addition for protein NMR experimentation.^28,29^ While liposomes are too large, micelles, bicelles, nanodiscs, and mmRMs are all amenable to protein NMR and have provided invaluable insight into protein-membrane interfacial interactions.^30^ However, these models have some drawbacks that may limit their practicality in protein-detected NMR fragment screening approaches.

Bicelles and nanodiscs are very large assemblies, making them less practical for screening due to the need for deuteration of the detergent and protein for even modestly sized proteins.^31^ Additionally, nanodiscs require preparation of a scaffold protein, an increased burden for screening which utilizes large numbers of samples.^32^ Micelles, while smaller, are generally constructed of artificial detergents that can distort the protein structure and limit sample stability.^5,33,34^ Alternatively, reverse micelles (RMs) have not only proven to be useful for a variety of biophysical protein NMR studies but they have been shown to be compatible with fragment screening of water-solubilized proteins.^35–38^ The recent development of mmRMs promises to extend the biophysical and fragment screening approaches developed in RMs to membrane-associated proteins.

The protein-lipid bilayer interface is a largely untapped avenue for drug discovery. To further complicate discovery efforts, libraries that are utilized for HTS screening campaigns tend to be populated with known inhibitors, natural products, and their analogs, which would limit the discovery of unknown classes of molecules.^16–18^ As membrane proteins become more prominent as drug and inhibitor targets, the need for novel classes of compounds is apparent. A fragment screening approach allows sampling of a large chemical space while avoiding the bias present in high-throughput libraries.^26,39^ Additionally, exploration of protein-membrane interface ligandability is worthwhile due to the recent attention to this mode of inhibitor binding.^6^ We used glutathione peroxidase 4 (GPx4) as our proof-of-principle system, a lipid hydroperoxidase that functions at cellular membranes to reduce lipid hydroperoxides.^40–42^ GPx4 is a highly sought after drug target due to its regulation of ferroptosis, an iron-dependent and non-apoptotic form of cell death^43–45^ which has shown promise in eliminating therapy-resistant cancer cells known as persister cells.^46–48^ GPx4 is a difficult target due to its lack of a deep active-site pocket and it functions as a lipid hydroperoxidase only in its membrane-bound state.^49^ The warheads currently used to target GPx4 are chloroacetamides, namely (1*S*,3*R*)-RSL3 (RSL3) and ML162, and masked electrophiles such as ML210. While masked electrophiles like nitrile oxides are prodrugs with some improved selectivity compared to the low stability and high promiscuity of the chloroacetamide probes,^49^ moving away from these covalent warheads altogether would avoid the disadvantages with this type of reactive electrophile.^50^ The footprint of GPx4 against its membrane is large and highly cationic^51,52^ precluding a strategy aimed at blocking membrane binding.^53,54^ In contrast, functional residues are localized to small, potentially ligandable regions of the protein within the membrane-interface, with the cationic patch implicated in lipid binding and the catalytic site for enzymatic activity.^51,52^ A non-covalent inhibition strategy may target this lipid binding region to prevent engagement with substrates. We sought to screen the functional, membrane bound state of GPx4 for fragment ligands, with the ultimate goal of assessing ligandability and unveiling potential building blocks for inhibitors that block lipid binding or catalysis in the membrane-protein interface.

To undertake a fragment screen of GPx4, we selected the most stable and suitable membrane model, which we found to be mmRMs comprised of a mixture of 1,2-dilinoleoyl-*sn*-glycero-3-phosphocholine (DLPC) and *N*-dodecyl phosphocholine (DPC) at concentrations of 37.5 mM each. A successful screen of the protein was conducted, identifying 9 fragment hits all with an apparent affinity less than 1 mM, demonstrating ligandability of the membrane-bound form of GPx4. The fragment hits spanned a range of partitioning coefficients and behavior. Importantly, they lacked significant binding to the water-solubilized form of GPx4, even at extreme concentrations. Finally, while the fragments weakly bind, secondary screening experiments were performed to demonstrate structure activity relationships (SAR) and the possibility to expand and enhance the small fragments towards a non-covalent inhibitor of GPx4. This approach promises to be a powerful tool in the fragment screening arsenal, has revealed fundamental properties of ligands that bind within protein-membrane interfaces, and has uncovered non-covalent fragments for GPx4 which may be starting points for a new class of inhibitors.

## Results and Discussion

### Membrane Model Selection

A variety of membrane models are available for the study of MPs and their relevant interactions, leaving us with the challenge of finding the best suited for fragment screening. Since protein NMR is one of the most preferred methods for fragment screening, we limited our membrane model selection to those that are compatible with this technique. Stability of the membrane model is important for fragment screening to ensure breakdown or expansion of the model is not misinterpreted as a false positive hit. To ensure solubility of fragments in aqueous buffer conditions, DMSO is commonly used, though we anticipated DMSO may interfere with established membrane models. To assess, we measured disturbance to the detergent assemblies upon introduction of 5% DMSO using Dynamic Light Scattering (DLS). Introduction of 5% DMSO to DPC micelles increased the size from 4.3 nm to 5.1 nm (**Figure S1a**), indicating incorporation of the co-solvent. DHPC:DMPC isotropic bicelles showed a second population is formed in the presence of 5% DMSO around 5.2 nm accounting for 20% of the total population, which indicates breakdown of the assembly (**Figure S1b**).^55^ While nanodiscs are excellent membrane models, their use in fragment screening by protein NMR is less practical due to their large size, requiring deuteration of lipid components, and the requirement to express and purify a scaffold protein in order to construct. For these reasons, we sought another alternative.

In contrast to micelles and bicelles, RMs are solubilized in a bulk alkane phase including a hexanol cosurfactant. The surfactants and cosurfactants form a spherical shell surrounding a nanoscale pool of water containing the protein.^37^ Recently developed mmRMs allow embedment of PMPs into the phosphocholine-rich inner shell as they would within a membrane.^28^ Hydrophilic small molecules are known to be highly soluble within the water core of RMs, while the alkane-hexanol mixed solvent enables solubilization of more hydrophobic small molecules.^35^ Together, the mixed solvent system promises to allow direct delivery of fragments to the mmRM system without the need for DMSO and with minimal perturbation of the membrane model. Fragments are generally screened in mixtures to increase throughput, so compatibility with fragment mixtures is essential. To confirm, we solubilized 10 mixtures of 10 fragments in separate mmRM samples, without DMSO. Of the 10 mixtures, only 1 mmRM sample had visually insoluble fragments. The mmRM size remains relatively constant with the addition of 10 fragments as observed by DLS (**Figure S1d**). A complementary experiment was performed with DPC micelles and the same 10 mixtures, without DMSO. In contrast with mmRMs, 4 out of 10 mixtures were observed to have insoluble aggregation in the DPC micelle samples. Size was assessed with DLS, showing similar variance as fragments with mmRMs (**Figure S1c**), yet the lower proportion of fragment solubilization and incompatibility with DMSO reduce the utility of micelles for fragment screening. In addition, mmRMs have other advantages over micelles and bicelles, such as enhanced protein stability compared to micelles, favorable tumbling properties compared to bicelles and nanodiscs, and simplicity of construction compared to nanodiscs.^30^ Together, these advantages and enhanced fragment compatibility led us to pursue mmRMs as a platform to fragment screen membrane bound PMPs, using GPx4 as our proof-of-principle target.

### Fragment screening within mmRMs

With mmRMs selected as our membrane model, we undertook a protein-observed, NMR-based fragment screen of GPx4. The screen was initially established by optimizing the encapsulation of GPx4 (**Figure 1a**). Once the mmRM conditions were optimized, a commercial, 1,911-member fragment library was screened, which contained fragments that are rule-of-three compliant,^21,23^ were filtered to avoid PAINS,^56^ and were selected to sample broad chemical diversity. Fragments were initially screened at 4:1 fragment to protein molar ratio in mixtures of 10 to enhance throughput. Use of mmRMs eliminated the need for DMSO to solubilize fragments. Encapsulation of GPx4 within mmRMs proceeded as previously reported with adjustments to the buffer.^28^ To deliver the fragments, the entire pre-constructed mmRM sample was transferred to the vial containing the pre-dried mixtures. Over 90% of the fragment mixtures were fully soluble in the mmRM, with the remaining 10% containing minor insoluble aggregates. Mixtures containing insoluble aggregates were nevertheless screened for fragment binding, with the assumption that fragments that could not be solubilized would not interfere and be misinterpreted as hits. No fragment hits were observed in samples containing insoluble aggregates, confirming that their presence did not inherently lead to false positives. The 191 mixtures of 10 fragments apiece were analyzed by ^15^N-HSQC experiments of GPx4. Any mixtures showing promising chemical shifts in the membrane-interacting surface were flagged as potentially containing a hit (**Figure 1b**). Fragment members of the 10 hit mixtures showing the most shifting were tested individually at a 4:1 ligand to protein ratio to reveal the identity of the hit within each mixture. To select hits from non-hits, we established chemical shift perturbation (CSP) cutoffs, with 7 out of the 15 observable cationic patch or catalytic site resonances mapped from previous experiments^28^ producing CSPs greater than 0.01 ppm. Using these CSP cutoffs, 14 individual fragments were identified as potential GPx4 binders. These fourteen fragments were characterized by titration to validate the hit and assess affinity.

**Figure 1.**
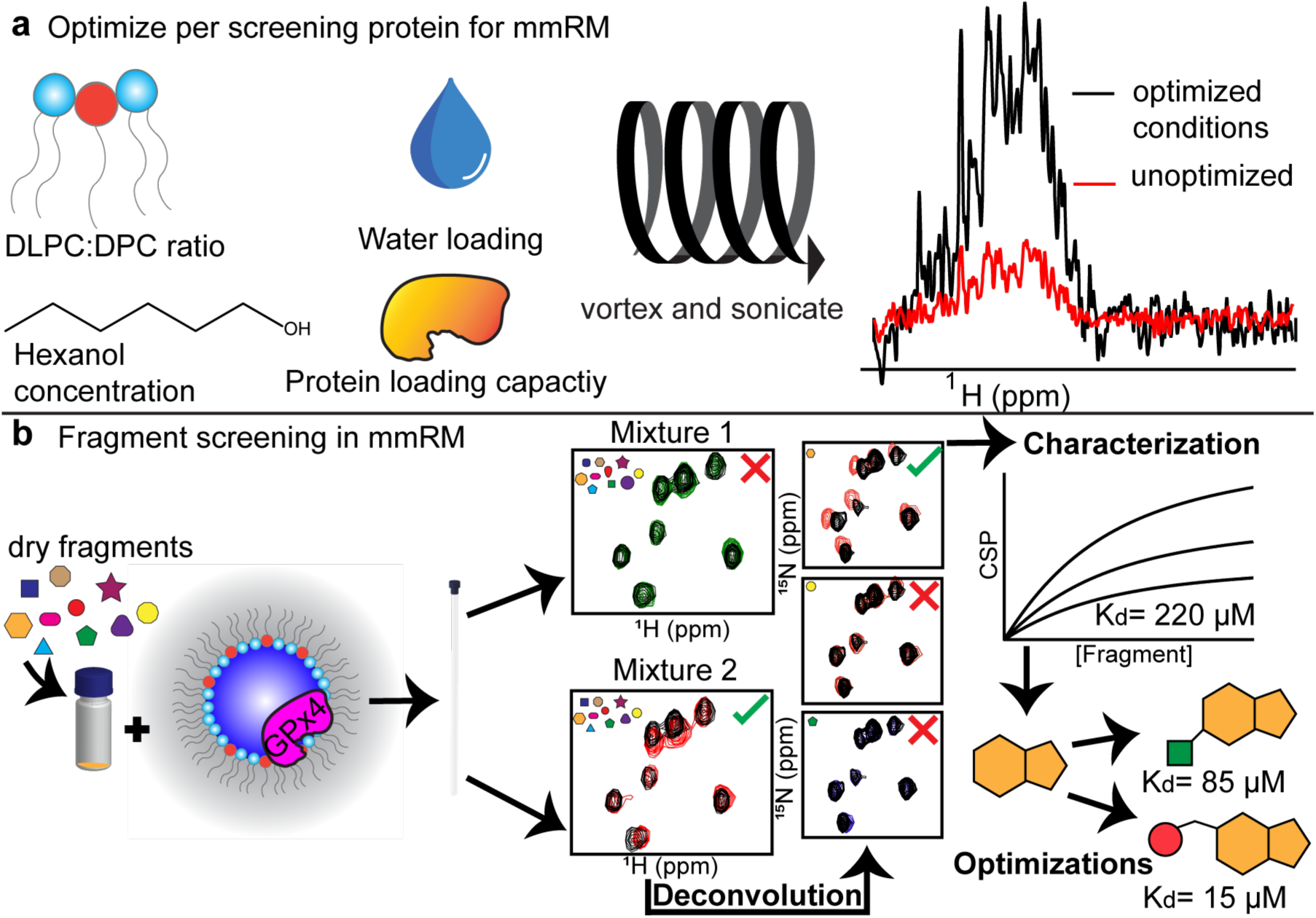
mmRM fragment screening workflow. **a)** Parameters such as DLPC:DPC ratio, water loading (water to surfactant molar ratio, W_0_), hexanol concentration, and protein concentration are optimized prior to the addition of fragment mixtures. **b)** Once optimized, fragment mixtures are dried and mixed with pre-constructed mmRMs with the protein of interest already encapsulated. Routine fragment mixture screening and deconvolutions by protein detected NMR is performed to isolate hits for the protein of interest. Characterization of hits was performed by NMR titrations to extract apparent K_d_s. Finally, optimizations of the fragments may lead to enhanced binders for the protein of interest.

### Fragment validation and binding affinity determination

To assess whether the fragment hits bound specifically to GPx4 and to calculate affinity, we performed NMR-based titrations. Chemical shift changes that approach saturation in a structurally localized manner target indicate specific binding.^27^ This allows true positive hits to be separated from false positives and weak or non-specific binders. Additionally, NMR based titrations allow for extraction of a binding affinity (K_d_). To calculate apparent K_d_ values, CSP cutoffs were used to isolate the resonances with most shifting using 1 standard deviation above the average CSP. All 14 fragments identified from the deconvolutions were titrated up to 400 µM or 1 mM if needed. Of the 14, 9 had apparent K_d_s less than 1 mM, and were identified as hits. Observed here is a 0.47% hit rate with best affinities in the µM range for the membrane-bound form of GPx4 and is consistent with ligandable and potentially druggable targets.^14,57^ The highest affinity fragment, fragment 1, had an apparent K_d_ of 105 µM ± 30 (**Figure 2a,b**). The remaining 8 fragment binding curves are shown in **Figure S2**. The remaining 5 fragments, 10-14, were ruled out as weaker (>∼1 mM) or non-specific binders. Apparent K_d_ values are summarized in **Table 1**. We note here that apparent K_d_ values reported in this study account for a total amount of fragment and protein within the mmRM sample and are not corrected for any potential concentrating effect due to localization of fragment to the mmRM surface or water phase.

**Figure 2.**
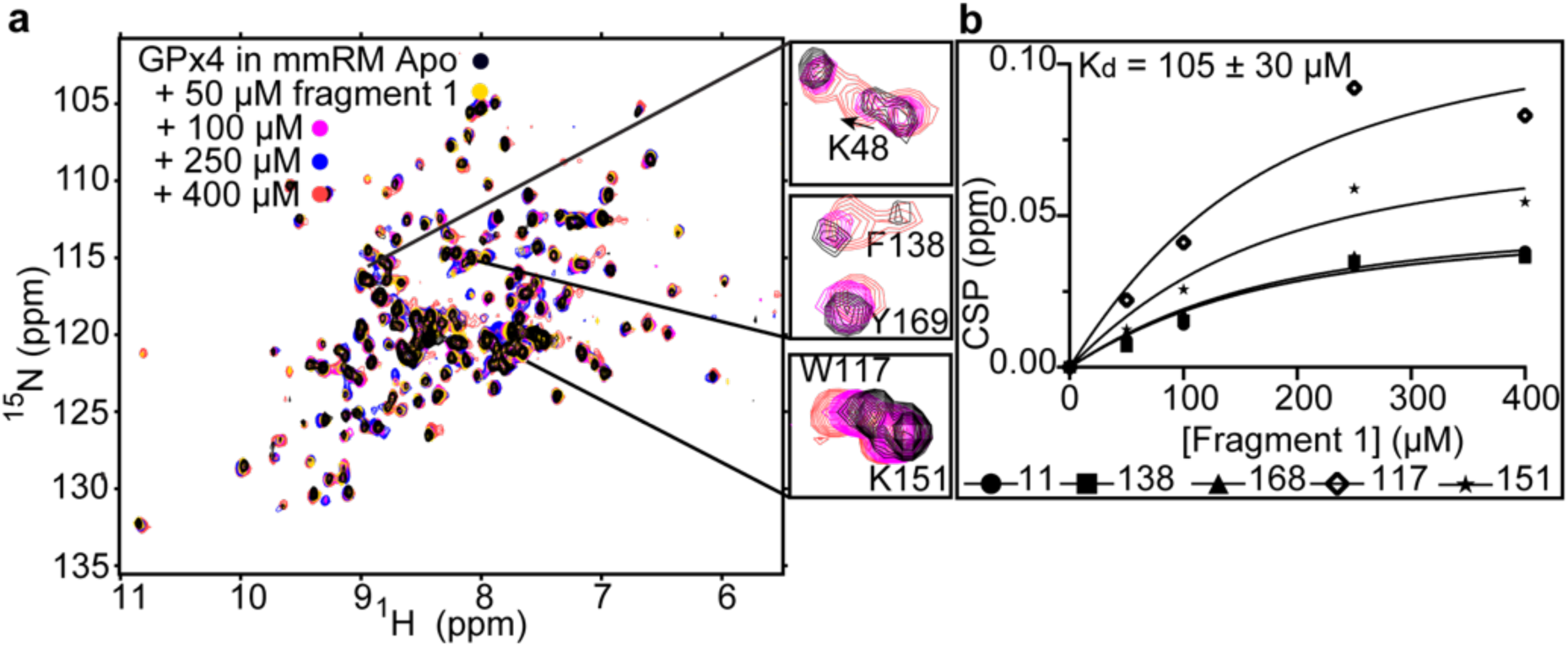
Titration of fragment 1 against GPx4 embedded in a mmRM. **a)** Overlay of [^15^N-^1^H] HSQC of GPx4 in the DLPC:DPC mmRM (black) with 50 (yellow), 100 (pink), 250 (blue), and 400 µM (red) of fragment 1. Zoomed panels show example resonances with chemical shift perturbations. For ease of visualization, only apo GPx4 along with 100 and 400 µM fragment 1 are shown in the zoom panels. **b)** Plot of representative chemical shift perturbations used for the apparent K_d_ extraction for fragment 1 against GPx4. Resonances used for K_d_ fitting were selected by identifying the top-sifting resonances at 400 µM fragment 1. These were resonances with a CSP at least 1 σ above the average CSP and corresponded to residue numbers 11, 117, 138, 151, and 168. These top-shifters were then fit to individual K_d_ curves and resonances with fits with a R^2^ < 0.85 was removed. The remaining resonances were then included in a global fit to determine the overall apparent K_d_.

**Table 1.**
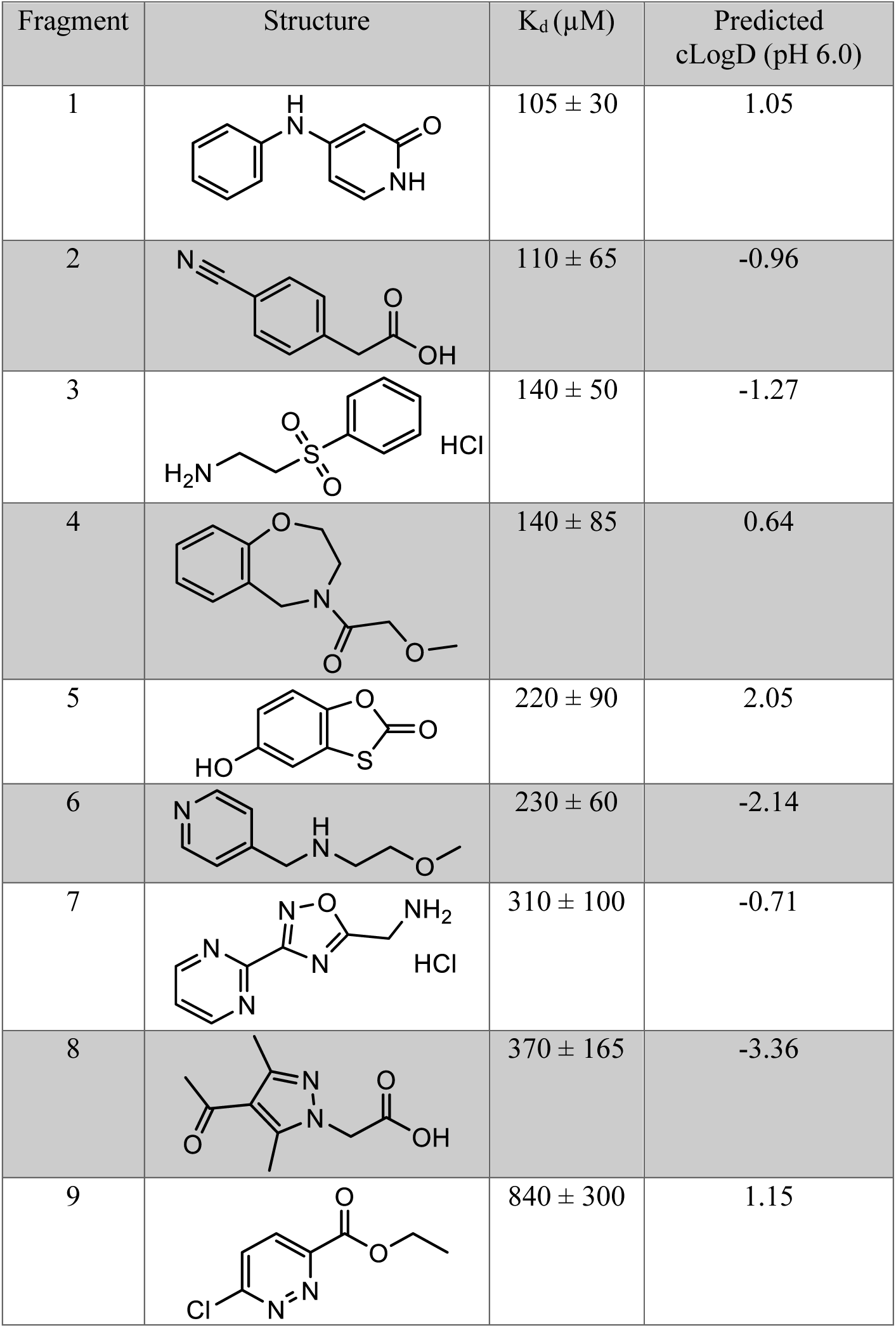
All fragments titrated against GPx4 in the mmRM with an apparent Kd less than 1 mM as assessed by NMR titrations.

| Fragment | Structure | $K_d$ (μM) | Predicted<br>cLogD (pH 6.0) |
| --- | --- | --- | --- |
| 1 | | $105 \pm 30$ | 1.05 |
| 2 | | $110 \pm 65$ | -0.96 |
| 3 | | $140 \pm 50$ | -1.27 |
| 4 | | $140 \pm 85$ | 0.64 |
| 5 | | $220 \pm 90$ | 2.05 |
| 6 | | $230 \pm 60$ | -2.14 |
| 7 | | $310 \pm 100$ | -0.71 |
| 8 | | $370 \pm 165$ | -3.36 |
| 9 | | $840 \pm 300$ | 1.15 |

Next, we sought to understand if fragment hits were solely bound to the membrane-embedded form of GPx4 or could also bind to the aqueous form. We performed aqueous binding experiments with 3 fragment hits with varying hydrophobicities. Of the fragment hits with K_d_ < 1 mM, the most hydrophobic (fragment 5, cLogP 2.05, 5-hydroxy-1,3-benzoxathiol-2-one), a hydrophilic fragment with the strongest affinity (fragment 6, cLogP −2.14, (2-methoxyethyl)(4-pyridinylmethyl)amine hydrochloride) were selected, along with a compound with an amphipathic structure and only modest hydrophobicity (fragment 1, cLogP 1.05, 4- (phenylamino)-1,2-dihydropyridin-2-one). Aqueous GPx4 was combined with 400 µM of either fragment 1, 5, or 6 to directly compare to the mmRM experiments. Comparing the CSPs in aqueous versus mmRM, we observed that the fragment hits do not bind strongly to the aqueous form of the protein (**Figure 3a-c**). This also suggests that a screen under aqueous conditions would not uncover these fragment hits. Since our fragments have variable degrees of solubility and partitioning, we realized a hydrophilic fragment like 6 may partition into the water core of the mmRM, essentially causing a concentration effect as observed previously.^35,36^ To rule this out, the concentration of the fragments was increased to 15 mM, which would be the effective concentration if all of the fragment partitioned completely into the RM water core. For fragments 1 and 6, neither produced significant CSPs (**Figure 3a,c**). Fragment 5 does produce some CSPs that could be detected by a soluble screen (**Figure 3b**). A very high concentration, 15 mM, was needed to reach CSPs comparable to the mmRM conditions in some resonances, which is much higher than the standard screening concentrations typically ∼100 – 400 µM. The 15 mM comparison of fragment 6 also demonstrates that observed CSPs are not purely the result of extremely weak binding due to a concentrating effect into the water component of the mmRM. DLS measurements were also collected to check for fragment aggregation in the aqueous conditions.^58^ At 400 µM, none of the fragments had any observable aggregation, but at 15 mM of fragment 1 and 5, there was a small population of large aggregating species, which was expected considering the cLogD values. Inspection of ^1^H-NMR spectra confirmed that the aggregation observed at high concentration of fragments 1 and 5 is a small portion of the overall fragment concentration and that most of the fragment is in solution. A lack of observed binding of these fragments to protein highlights that they do not bind to the water-solubilized state of GPx4, with binding of the hydrophobic and amphipathic fragments driven to the membrane interface. This is possibly due to a conformational change upon membrane binding, which is suggested by the very large chemical shift changes in the interfacial region when GPx4 binds to membrane models.^28,29,52^

**Figure 3.**
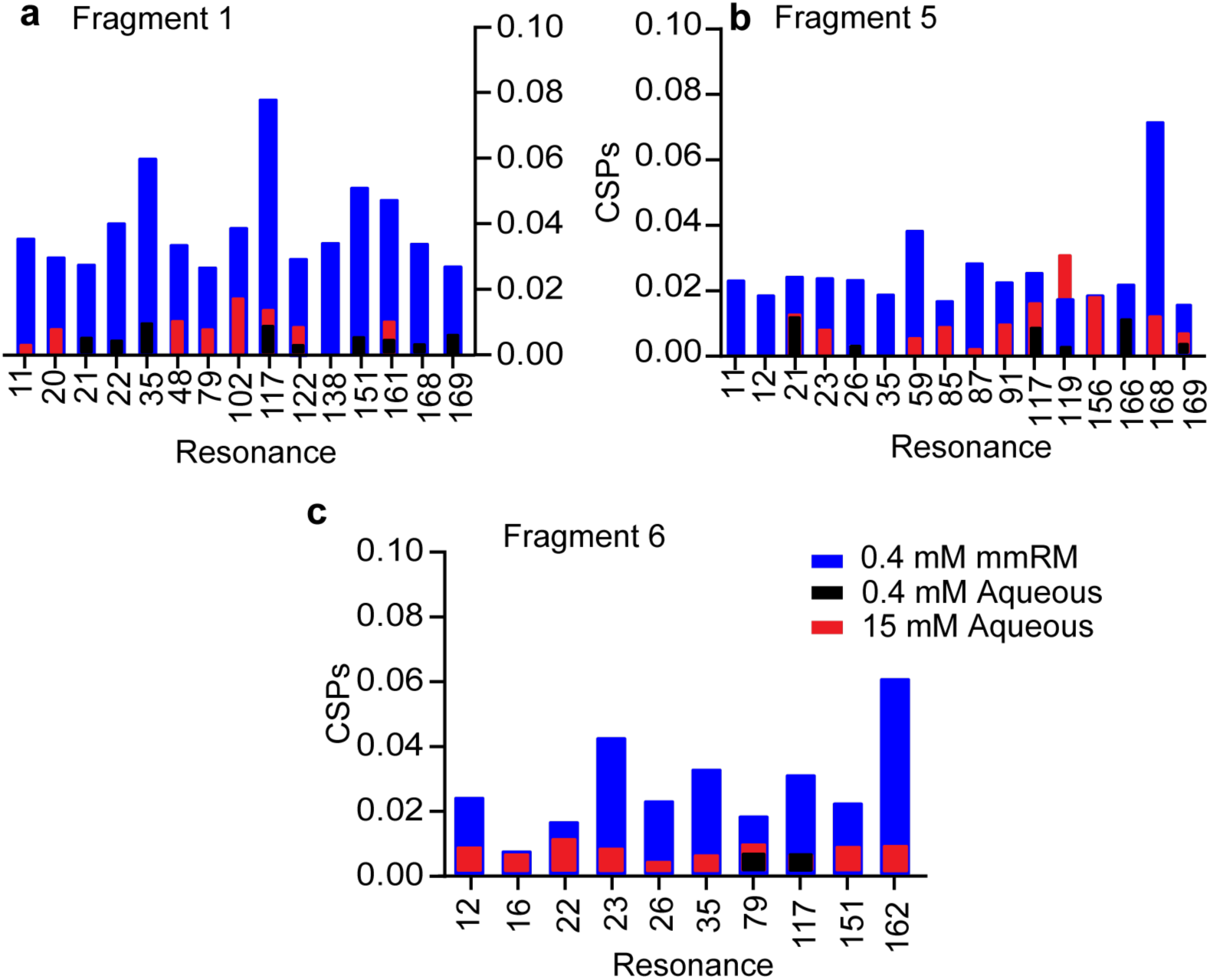
Comparison of hit binding to membrane-embedded versus aqueous GPx4 for **a)** fragment 1, **b)** fragment 5, and **c)** fragment 6. [^15^N-^1^H] HSQC experiments were conducted with 400 µM of fragment 1, 5, or 6. The resonances that produced the greatest CSPs (at least 1σ above the average) for the mmRM experiment (blue) are compared to the corresponding aqueous data. The identical experiment was performed in bulk aqueous conditions (black). 15 mM experiments were also conducted (red) to test for very weak binding.

To visualize the fragment binding sites, the top shifting resonances in the membrane interface by CSP analysis for fragments 1, 5, and 6 were mapped onto the crystal structure of GPx4. Both fragments 1 and 5 have several shifting resonances in the membrane interface, indicating the fragments are able to partition into the mmRM membrane model to bind the protein (**Figure 4a,b, S3a,b**). We note that due to some line-broadening in the membrane interface of GPx4, not all interface residues are observable. For fragment 1, the significantly shifting residues, A11, K48, G79, W117, I122, F138, K151, V161, H168, and Y169, were within or neighboring the cationic and catalytic sites of GPx4 and the membrane interface.^28^ For fragment 5, shifting residues were similarly A11, R12, W117, W119, C148, M156, L166, and Y169. Fragment 6 seems to have less interaction with the protein at the membrane interface, which could potentially be a result of the overall hydrophilicity of the fragment in comparison to fragments 1 and 5 and binding to the water-exposed allosteric site (**Figure 4c, S3c**).^59^ Some membrane interface residues did display shifting with G79, W117, G126, K151, and I162 having the most. Regardless, presence of the membrane model was necessary for the fragment interaction as observed in **Figure 3c**. A better understanding of fragment partitioning capabilities could lead to enhanced GPx4 targeting, and therefore fragment optimization and elaboration.

**Figure 4.**
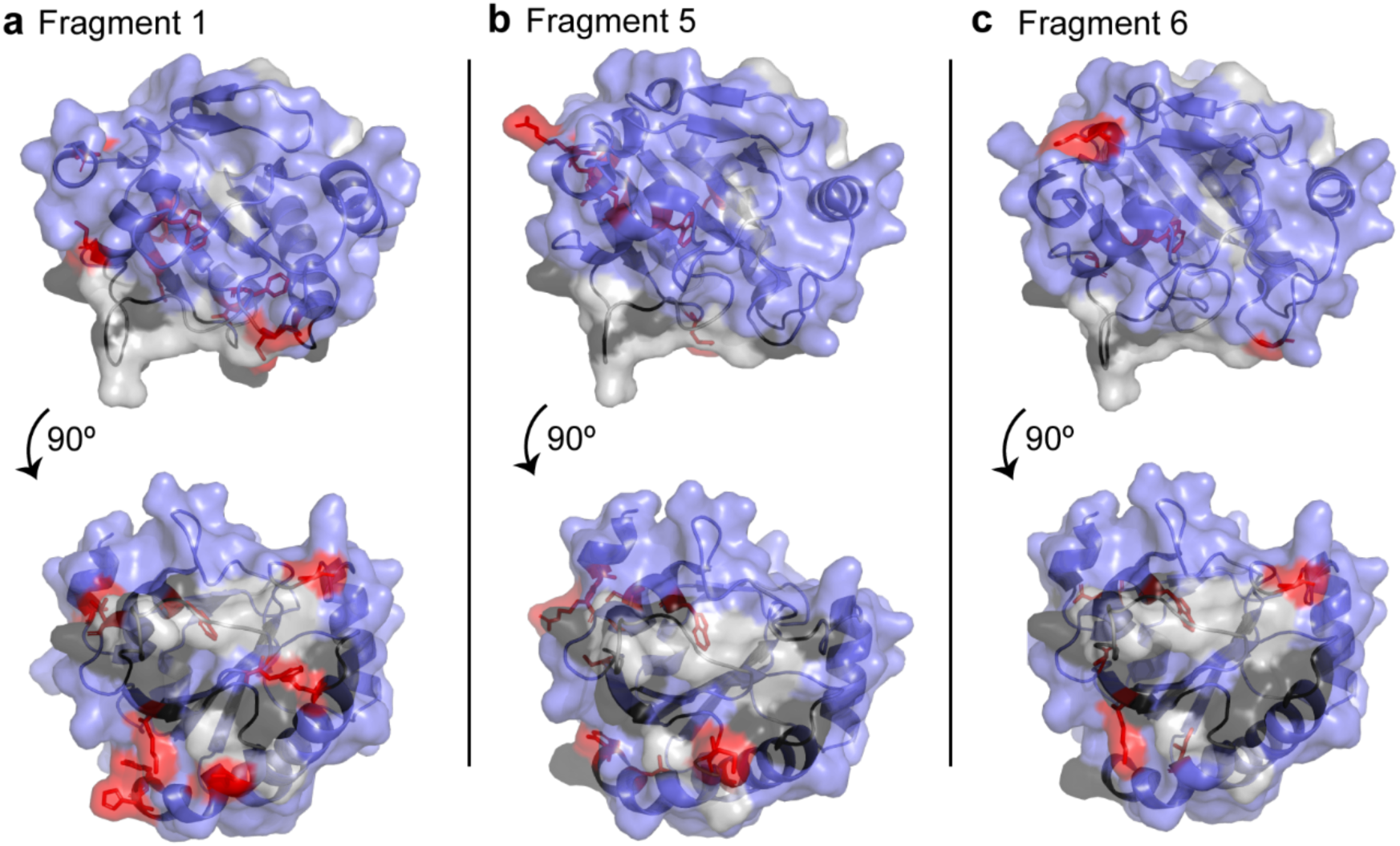
Mapped chemical shift perturbations for the membrane interacting residues of GPx4 with **a)** fragment 1**, b)** fragment 5, and **c)** fragment 6 are displayed with red sticks. Residues that are unassigned in the mmRM are displayed in light gray, residues with little or no shifting are shown in blue, and other membrane interacting residues are in black.

### Partitioning properties of fragments in mmRMs

Interestingly, the 9 validated hits had varying predicted partitioning values (cLogD, pH = 6.0) between −3.36 to 2.05, spanning from hydrophobic to hydrophilic **(Table 1).** We sought to understand the potential membrane partitioning of these fragments by evaluating their behavior in mmRMs, again using fragments 1, 5, and 6 as our benchmarks. To evaluate fragment partitioning in the mmRM system, 2D ^1^H-^1^H NOESY experiments were performed, which report on spatial proximity of intra- and intermolecular nucleii. Initial experiments demonstrated 5 mM fragment is needed to clearly observe intermolecular NOEs. The mmRMs were constructed from deuterated pentane and hexanol to reduce solvent signal and associated NOESY artifacts, which were still present, but NOESY measurements were possible.^60^ Our moderately hydrophobic fragment, fragment 1 with a logD of ∼ 1 shows clear NOE crosspeaks to the surfactant shell, in particular to the surfactant tails and headgroup, and potentially with water (**Figure 5a**). Fragment 5 was the most hydrophobic fragment hit from this screen with a predicted logD of ∼2. The 2D NOESY reveals the majority of the fragment must reside in the alkane phase of the reverse micelle since there does not appear to be any relevant crosspeaks between the fragment and the water or the surfactant shell (**Figure 5b**). A lack of NOEs to pentane is due to the necessity of using deuterated pentane for artifact reduction and attempts to directly observe solvent NOEs failed. The most hydrophilic fragment, fragment 6 with a predicted logD of about −2, is expected to predominantly have intermolecular interactions with the water phase of the mmRM (**Figure 5c**). However, interference from the water peak introduced streaking and precluded the visualization of clear crosspeaks.

**Figure 5.**
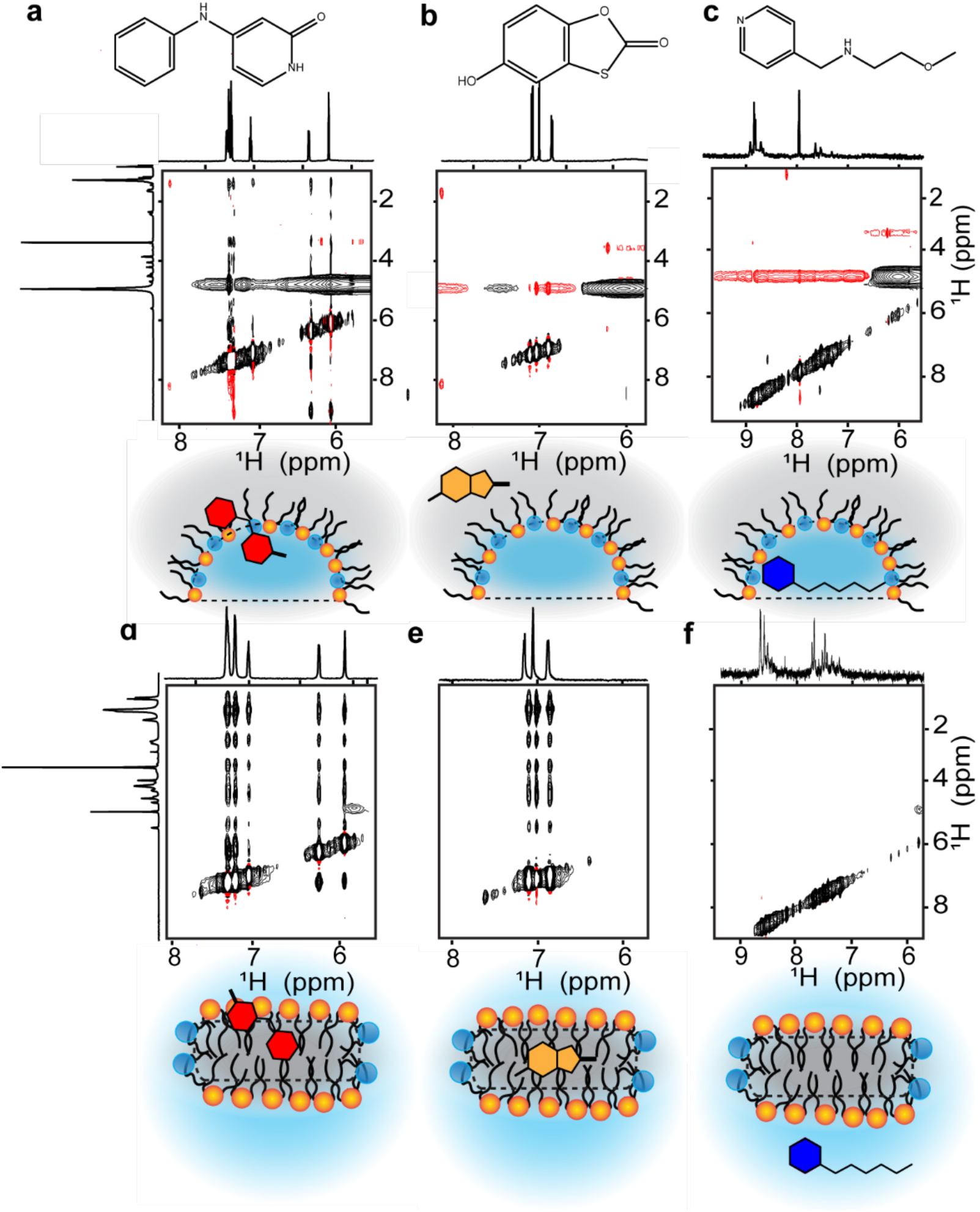
Zooms of [^1^H-^1^H] NOESYs of the 3 fragments show variable partitioning in the mmRM. **a)** The most amphipathic fragment, fragment 1, shows clear interaction with the surfactant shell of the mmRM as well as with water. **b)** Fragment 5, the most hydrophobic fragment, does not show any clear NOEs, indicating the fragment may be entirely residing in the pentane phase in the absence of GPx4. **c)** The most hydrophilic fragment, fragment 6, does not have any NOEs to the surfactant shell and may be fully residing in the water core of the mmRM. Positive contours are shown in black and negative contours are shown in red. Zooms of [^1^H-^1^H] NOESYs of the 3 fragments in a DMPC:DHPC bicelle to further probe partitioning. All bicelle samples were in D_2_O to reduce the water signal to better observe NOEs. **d)** Fragment 1 has interactions with the surfactant shell of the bicelle similarly to the mmRM, but **e)** fragment 5 is within the bicelle core **f)** Fragment 6, now in a completely D_2_O phase, still does not have any observable NOEs. Cartoon representations have been included for partitioning visualization.

To confirm partitioning behavior, we tested fragments in DHPC:DMPC bicelles, which uses a bulk aqueous solvent and the hydrophobic phase is confined to the bicelle interior. To successfully collect NOESY data, the sample solvent was composed of 100% D_2_O. As expected, fragment 1 partitions to the bicelle, similarly to the mmRM (**Figure 5d**). NOEs between fragment 5 and all components of the surfactants, indicate full partitioning into the hydrophobic core of the bicelles (**Figure 5e).** For fragment 6, no NOE crosspeaks to bicelles surfactants were observed (**Figure 5f**) and no water cross peaks were observed due to necessity of 100% D_2_O to reduce artifacts. This result confirms fragment 6 prefers to reside in the water core of the mmRM and the aqueous solvent in the bicelle system. Together, the results highlight the partitioning behavior of fragments that bind to GPx4. Hydrophilic, hydrophobic, or fragments that partition to the surfactant shell are all capable of binding to the membrane-bound state of GPx4 and may be detected using this method.

### Enhanced affinity through fragment optimization

To investigate whether or not these fragments could be built upon and progress to higher affinity binders, and potentially inhibitors, a secondary screening approach was used to find analogs of the original fragments. A fragment growing approach was followed due to the high success rate of improving fragment hits. ^61,62^ Fragment growing improves binding affinity, or other properties, with the addition of different substitutions and/or expansions.^23^ Commercially available analogs of fragments 1 and 5 were found by searching the ChemBridge (San Diego, CA) catalog for analogs with a 60% similarity for fragment 1 and a 70% similarity for fragment 5. A subset of analogs for each fragment was purchased and tested. These two fragments were the main focus for fragment progression due to their observed ability to partition into bilayers and bind to inhibitory regions of membrane-bound GPx4. Fragment 6 was not investigated further since it resides mainly in the water core of the mmRM model and, while interesting, it falls outside of our focus for this study. While all five selected fragment 1 analogs produced reasonable CSPs in the membrane interface of GPx4, none produced titration curves that indicated enhanced binding (**Table S1**). This result led us to believe commercially available analogs alone would not lead to the productive development of this specific fragment. Alternatively, for fragment 5, a clear progression of binding affinity and structure optimization became apparent from titrations with compounds containing a phenyl and additional substitutions around the ring, namely mono- or dichloro or methyl substitutions in the para- or meta-positions (**Table 2)**. Interestingly, addition of the phenyl ring alone reduced affinity and the presence of at least one chloride was necessary to enhance the affinity. Of the seven analogs of fragment 5 selected for titrations, five fit to apparent K_d_s lower than the original fragment, ranging from 15-85 µM. Additionally, all of the analogs with an enhanced affinity were more hydrophobic than the original fragment. Meaning, as a whole, these analogs would also partition into membranes efficiently. Addition of hydrophobic groups is a suggested strategy to enhance affinity of inhibitors to membrane-proteins,^6^ which is reflected here. To validate this conclusion, analog 5.5, which has the highest cLogP of 4.90 with a high affinity of ∼30 µM, was selected for experimentation in DPC micelles and again in bulk aqueous conditions. The parallel experiment was also completed in mmRMs (**Figure S6a,b**), with 250 μM analog 5.5. These experiments will reveal whether analog 5.5 can target the protein in a membrane model but will not be an effective binder in the absence of the membrane. As expected, reasonable CSPs were observed in the DPC micelle experiment in the membrane interacting interface of the protein, while the comparative aqueous experiment showed little to no shifting that would indicate the analog was binding to the protein (**Figure 6a,b**). DPC micelles have somewhat less line-broadening in the membrane interface compared to the mmRM, allowing a more complete structural map of analog binding.

**Figure 6.**
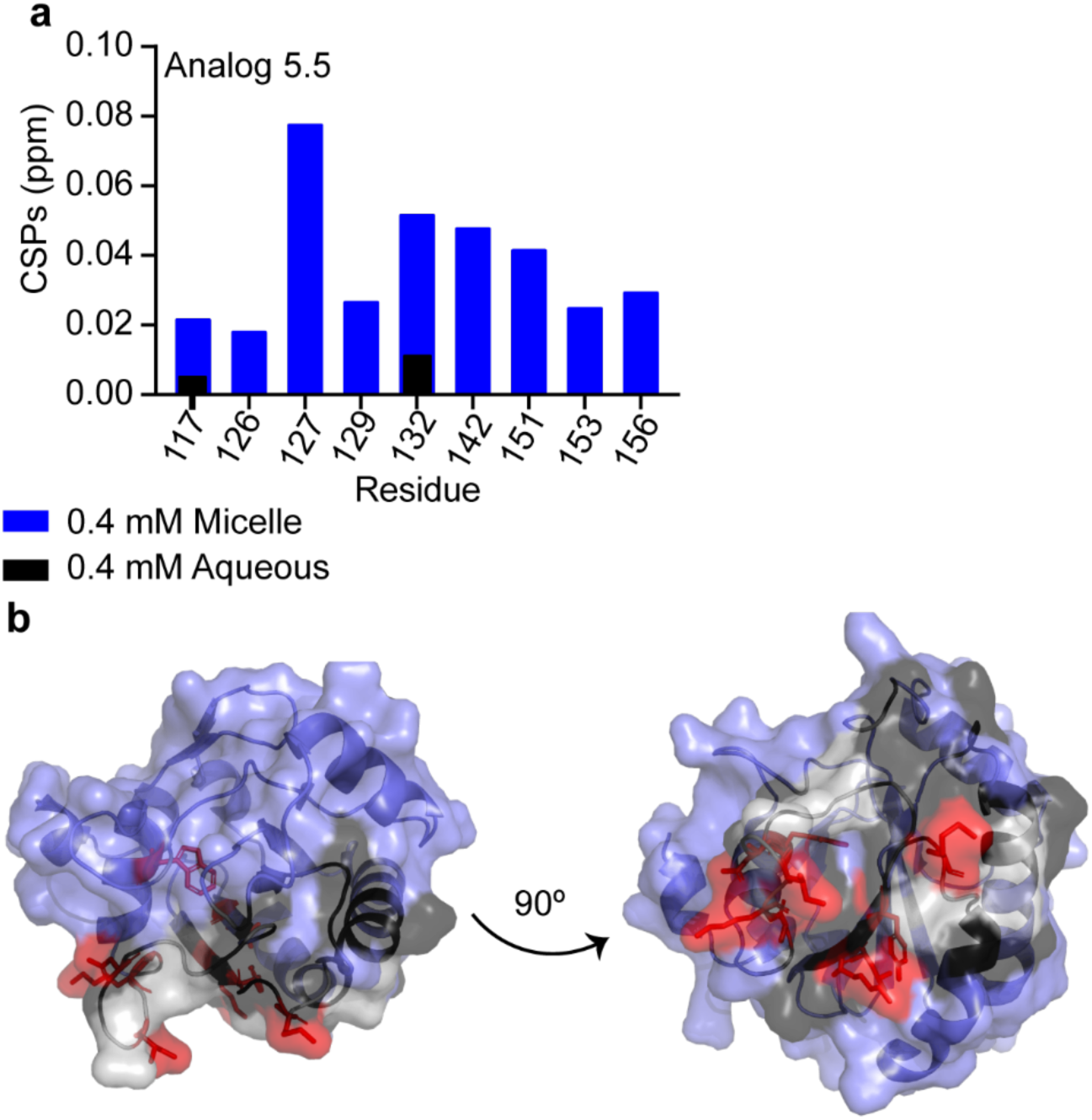
Analog 5.5 in DPC micelles. a) Top-shifting resonances from [^1^H-^15^N] HSQC experiments were identified by calculating the CSP between GPx4 bound to DPC micelles with 0.4% DMSO and GPx4 bound to DPC micelles with 400 µM analog 5.5 an 0.4% DMSO. CSPs at least 1σ above of the average were isolated as the top-shifters (blue bars). CSPs from the comparable aqueous experiment that consisted of apo GPx4 with 1.5% DMSO and GPx4 with 400 µM analog 5.5 an 1.5% DMSO are shown in black bars. The increased DMSO was required for full solubilization in these conditions. **b)** The top-shifting resonances were then mapped on a crystal structure of GPx4 with the membrane interface in DPC micelles shown in black, residues with little or no shifting are shown in blue, missing resonances in grey, and analog interacting residues shown with red sticks.

**Table 2.**
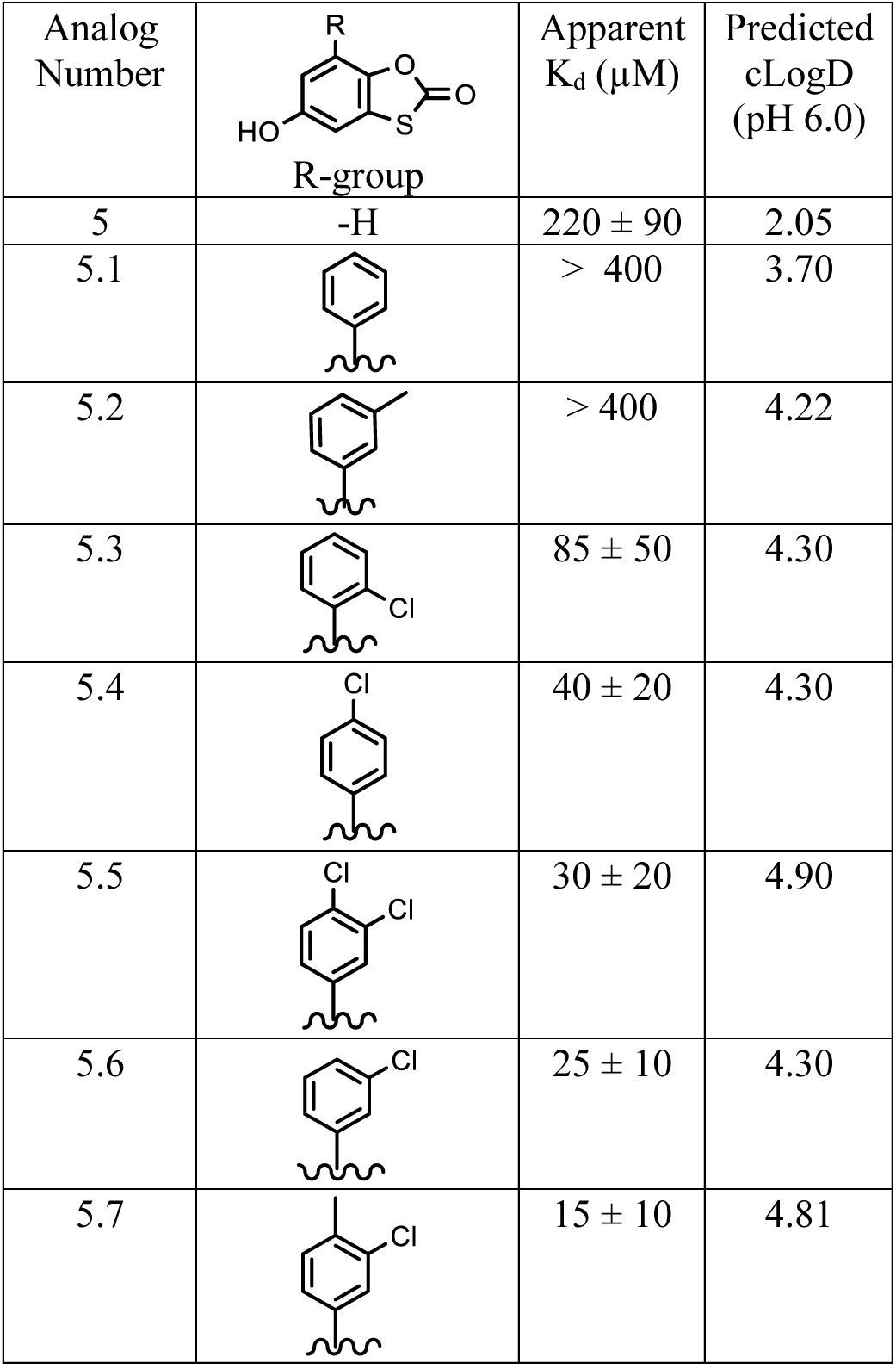
Apparent K_d_s and predicted cLogDs of titrated analogs of fragment 5.

| Analog Number | R-group | Apparent $K_d$ ($\mu\text{M}$) | Predicted cLogD (pH 6.0) |
| --- | --- | --- | --- |
| 5 | -H | $220 \pm 90$ | 2.05 |
| 5.1 | | $> 400$ | 3.70 |
| 5.2 | | $> 400$ | 4.22 |
| 5.3 | | $85 \pm 50$ | 4.30 |
| 5.4 | | $40 \pm 20$ | 4.30 |
| 5.5 | | $30 \pm 20$ | 4.90 |
| 5.6 | | $25 \pm 10$ | 4.30 |
| 5.7 | | $15 \pm 10$ | 4.81 |

While not ideal for screening conditions, other membrane models such as DPC micelles can be useful for characterizations of some individual hits. Altogether, these results show the original fragments have the potential to be developed into higher-affinity binders of GPx4. This approach could lead to a non-covalent inhibitor for GPx4 or other similar PMP targets that have thus far proven elusive for non-covalent small-molecule inhibition.

## Conclusions

Presented here is a new methodology that enables screening of small molecules and fragments for PMPs while the protein is engaged to the membrane. Use of mmRMs proves to eliminate some of the disadvantages of using other membrane models for fragment screening of membrane-engaged targets. This methodology allowed for the discovery of 9 fragment ligands and highlights the ligandability of GPx4 in its membrane bound state, contrasting the water-solubilized state which is considered unligandable by non-covalent inhibitors. These fragments may be used as building blocks for development of non-covalent small-molecule inhibitors for GPx4, which have not yet been reported. The fragments span a broad range of partitioning coefficients, an advantage of using the mmRM model and its bulk-nonpolar environment.

Partitioning properties were investigated using [^1^H-^1^H] NOESY experiments to reveal that regardless of where the fragments prefer residing, they can target and bind GPx4. Additionally, the potential to enhance and build off of these fragments was shown through a structure activity relationship study. A secondary screen of fragment 5 analogs revealed a series of compounds with higher affinity, revealing a potential pathway towards developing inhibitors.

Importantly, these fragment hits would not have been discovered in the absence of a membrane model, demonstrating that GPx4 is more tractable in its membrane-bound, active form. Regardless of the platform, mmRM or micelle, direct targeting of the protein was not observed unless GPx4 was engaged with a membrane model. Our results suggest the presence of a small-molecule binding site may depend on membrane engagement.^6^ The binding site that the fragments engage with may be due to a confirmational change upon membrane binding, or perhaps the local environment of the protein surface that is solvated by lipids may be uniquely poised to bind these fragments. Further study is needed to reveal details.

While GPx4 is a PMP, there may be utility for this screening technology for transmembrane proteins to target the non-solvated parts of the proteins. RMs are known to house transmembrane proteins efficiently, allowing this as a potential avenue for screening these difficult targets.^38,63^ This method may be applied to MPs broadly, and can be added to the repertoire of FBDD methodologies used to efficiently probe chemical space^26,27^, take into account inhibitor-lipid interactions,^7^ assess ligandability,^14^ and initiate drug discovery endeavors with a better consideration of membrane partitioning properties.^6^ Often, drug design and development relies on the assumption that there are only nonspecific interactions within the membrane, making them unexploitable.^1^ There has been an increasing number of membrane protein structures with binding sites displaying displacement of membrane lipids upon ligand binding, indicating tractability in the membrane interface.^6,64,65^ All of these considerations point to the pressing need to expand the use of membrane based screening platforms to ensure the largest set of tools are available for these challenging targets.

## Methods and Materials

Methods for micelle, bicelle, and mmRM construction, DLS measurements, and GPx4 protein production were performed as previously reported.^28,52^ Details can be found in **Supplementary Information.**

### Protein ^15^N-^1^H HSQC

NMR samples were prepared with ^15^N-isotopically labeled protein with 10% v/v of *d*-pentane as the lock solvent (Sigma-Aldrich, St. Louis, MO). All NMR spectra were collected at 25 °C on a 700 MHz on Bruker Avance III instrument. All NMR spectra were processed using NMRPipe^66^ and analyzed using NMRFAM-Sparky.^67^ Chemical shifts for GPx4 (BMRB 50955) were assigned from those previously published.^52^ Chemical shift perturbations from ^15^N-HSQC spectra were calculated using the following formula:

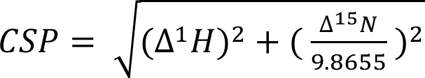

where Δ^1^H and Δ^15^N are the changes in ^1^H and ^15^N chemical shifts.

### Fragment delivery and screening

A custom subset of 1,911 members of the LifeChemicals high-solubility fragment library was used for this study. The subset members were selected to prioritize chemical diversity while still adhering to fragment rule-of-three metrics, to include only fragments with measured solubility confirmed at 1 mM in PBS, and to avoid PAINS compounds.^21,56^ To conduct the fragment screening, mixtures of 10 fragments were pipet into individual vials for a total concentration of 400 µM for each fragment. Samples of mmRM encapsulated GPx4 were transferred to the vials of pre-dried fragments, an additional 50 µM of hexanol was added, and subsequent vortexing and water bath sonication ensured the dried fragment was incorporated into the mmRM. The ∼190 mixtures of 10 fragments apiece were analyzed by ^15^N-HSQC experiments of GPx4. Spectra of each fragment mixture was compared to ^15^N-HSQCs of encapsulated GPx4 containing no fragment. Any mixture showing promising chemical shifts in the membrane-interacting surface was flagged as potentially containing a hit. From there, the best 15 mixtures were prioritized into 3 groups of 5 mixtures, which we termed high-priority, medium-priority, and low-priority. The five highest priority mixtures produced spectra with significant shifting with multiple resonances in the interface. Medium priority mixtures had less overall shifting, but still contained multiple resonances of interest while low priority mixtures had minimal shifting with 1-2 resonances of interest. It is important to note that this ranking system, while incorporating certain benchmarks, was qualitative and performed by eye. Moving forward, for the sake of efficiency, only the 10 high and medium priority mixtures were progressed forward to the deconvolution stage.

Fragment members of the top 10 hit mixtures were tested individually to reveal the identity of the hit within each mixture. 400 µM of each fragment was encapsulated with GPx4 and compared against the ^15^N-HSQC spectrum of GPx4 in the mmRM without any fragment. CSP cutoffs were established to isolate the hits from the non-hits with a minimum of 7 out of the 15 cationic patch or catalytic site resonances producing CSPs greater than 0.01 ppm denoting a hit. 14 fragments were identified as GPx4 binders using the CSP cutoffs.

### Fragment and Analog Titrations and Aqueous Screen

The 14 fragments isolated from deconvolutions were either purchased from LifeChemicals (Fragment 1, 4, 6, and 11, and 12-14), Combi-Blocks (2, 3, 5, 9, and 10), or ChemBridge Hit2Lead (7 and 8). All fragment 1 and 5 analogs were also purchased from ChemBridge Hit2Lead. Fragments and analogs purchased from LifeChemicals and Hit2Lead are 90+% pure while the fragments purchased from Combi-Blocks ranged from 95-97% pure. The day prior to the titration, a double sample of encapsulated GPx4 was made before splitting up into two separate vials. Half of the sample was added to the vial containing pre-dried fragment corresponding to 400 µM or 1 mM. The following day, the sample lacking fragment was used as the zero point of the titration and was compared to the high point, 400 µM or 1 mM of fragment, prior to the start of the titration to evaluate if significant CSPs were observed in the membrane interface region. If they were, then the two samples were mixed to produce varying concentrations and to measure a titration curve. The top-shifting resonances were determined by calculating the average CSP at 400 µM or 1 mM and 1 and 2 standard deviations (σ) above that average. The CSP for the resonance at each fragment concentration was extracted and fit to a curve using an affinity equation which accounts for ligand depletion:

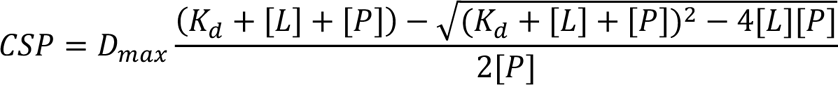

Any resonance producing a curve with an R^2^ below 0.85 was removed and the remaining resonances were then used to fit a group apparent K_d_ curve for the fragment hit.

Fragments 1, 5, and 6 were selected to perform proof-of-concept experiments in bulk aqueous conditions. 400 µM and 15 mM aliquots of fragment were vacuum dried overnight to remove excess DMSO. An apo GPx4 reference was collected with 1% DMSO or 5% DMSO to ensure CSP calculations took into account DMSO needed for fragment solubilization. 100 µM GPx4 was added to the dried fragments and DMSO was added to a total of 1% for the 400 µM and 5% for the 15 mM fragment experiments. These percentages were the lowest amount needed to solubilize the fragments in solution. Prior to experimentation, the pH of the sample was checked to ensure no change was observed upon the addition of the fragments. However, the 100 mM Bis-Tris pH 6.0 ensured the pH was not altered. ^15^N-HSQC experiments were collected and CSPs were analyzed and compared to the top-shifting resonances determined from the mmRM experiments. DLS measurements were collected to assess fragment aggregation in the aqueous conditions using a Malvern Zetasizer Nano-S instrument. At 400 µM, the fragments were combined in buffer with 1% DMSO and at 15 mM fragment 5% DMSO was used to enhance solubility.

### Fragment Partitioning by ^1^H-^1^H NOESY

Partitioning coefficients for fragment hits and analogs were calculated using Marvin LogD calculator (ChemAxon) and were used to prioritize fragments for 2D NOESYs. mmRMs were premade as previously described and loaded with 5 mM of dried fragment 1, 5, or 6 before shaking overnight. A high concentration of fragment is needed for these experiments to observe the intermolecular NOEs between the fragments and the membrane model components. For the DHPC:DMPC samples, the q = 1.0 bicelles were constructed routinely,^55^ as described in the supplemental information, with an additional step to ensure the complete removal of water before experimentation. An aliquot of buffer (100 mM Bis-Tris pH 6.0, 100 mM NaCl, and 10 mM DTT) was frozen and freeze dried overnight to remove residual water. The dried buffer was then resuspended in 100% D_2_O, mixed, and frozen before another round of freeze-drying. The components were then resuspended in 100% D_2_O again, before incorporating into the DHPC fraction of the bicelle. The DHPC solution was then mixed with dried DMPC and bicelle formation proceeded. NOSEY experimental design was based off of previous work.^60^ All NMR spectra were collected at 25 °C on a 700 MHz on Bruker Avance III instrument. The 2D data was collected with a 90⁰ pulse width between 10.2 and 11.0 µs, which was optimized prior to each experiment. 128 complex increments were collected in State-TPPI mode with 8 scans per increment, 2 s recycle time, and 250 ms mixing time. All spectral data was processed with NMRPipe^66^ and analyzed with NMRFAM-Sparky.^67^

## Author Contributions

C.L.L. performed all experiments. C.L.L. and B.F. designed experiments, analyzed and interpreted data, and wrote the manuscript. All authors have given approval to the final version of the manuscript.

## Notes

The authors declare no competing financial interest.

## Acknowledgments

We thank Dr. Santiago Lima, Sara Walters, and Jake Breeden for helpful discussions and technical assistance. Research reported here was supported by the National Institute of General Medical Sciences of the National Institutes of Health under Award Number R35GM147221. The content is solely the responsibility of the authors and does not necessarily represent the official views of the National Institutes of Health.

## References

(1) Yin, H.; Flynn, A. D. Drugging Membrane Protein Interactions. Annu Rev Biomed Eng 2016, 18, 51–76.

(2) Grauffel, C.; Yang, B.; He, T.; Roberts, M. F.; Gershenson, A.; Reuter, N. Cation− π Interactions as Lipid-Specific Anchors for Phosphatidylinositol-Specific Phospholipase C. J Am Chem Soc 2013, 135 (15), 5740–5750.

(3) Moravcevic, K.; Oxley, C. L.; Lemmon, M. A. Conditional Peripheral Membrane Proteins: Facing up to Limited Specificity. Structure 2012, 20 (1), 15–27.

(4) Boes, D. M.; Godoy-Hernandez, A.; McMillan, D. G. G. Peripheral Membrane Proteins: Promising Therapeutic Targets across Domains of Life. Membranes (Basel) 2021, 11 (5), 346.

(5) Gulezian, E.; Crivello, C.; Bednenko, J.; Zafra, C.; Zhang, Y.; Colussi, P.; Hussain, S. Membrane Protein Production and Formulation for Drug Discovery. Trends Pharmacol Sci 2021, 42 (8), 657– 674.

(6) Payandeh, J.; Volgraf, M. Ligand Binding at the Protein–Lipid Interface: Strategic Considerations for Drug Design. Nat Rev Drug Discov 2021, 20 (9), 710–722.

(7) Peetla, C.; Stine, A.; Labhasetwar, V. Biophysical Interactions with Model Lipid Membranes: Applications in Drug Discovery and Drug Delivery. Mol Pharm 2009, 6 (5), 1264–1276.

(8) Fernandes, E.; Cardoso, V. F.; Lanceros-Méndez, S.; Lúcio, M. Lipid Microfluidic Biomimetic Models for Drug Screening: A Comprehensive Review. Adv Funct Mater 2024, 2315166.

(9) Li, H.; Zhao, T.; Sun, Z. Analytical Techniques and Methods for Study of Drug-Lipid Membrane Interactions. Rev Anal Chem 2018, 37 (1), 20170012.

(10) Tubiana, T.; Sillitoe, I.; Orengo, C.; Reuter, N. Dissecting Peripheral Protein-Membrane Interfaces. PLoS Comput Biol 2022, 18 (12), e1010346.

(11) Coleman, N.; Rodon, J. Taking Aim at the Undruggable. American Society of Clinical Oncology Educational Book 2021, 41, e145–e152.

(12) Brown, D.; Superti-Furga, G. Rediscovering the Sweet Spot in Drug Discovery. Drug Discov Today 2003, 8 (23), 1067–1077.

(13) Cheng, A. C.; Coleman, R. G.; Smyth, K. T.; Cao, Q.; Soulard, P.; Caffrey, D. R.; Salzberg, A. C.; Huang, E. S. Structure-Based Maximal Affinity Model Predicts Small-Molecule Druggability. Nat Biotechnol 2007, 25 (1), 71–75.

(14) Edfeldt, F. N. B.; Folmer, R. H. A.; Breeze, A. L. Fragment Screening to Predict Druggability (Ligandability) and Lead Discovery Success. Drug Discov Today 2011, 16 (7–8), 284–287.

(15) Surade, S.; Blundell, T. L. Structural Biology and Drug Discovery of Difficult Targets: The Limits of Ligandability. Chem Biol 2012, 19 (1), 42–50.

(16) Hert, J.; Irwin, J. J.; Laggner, C.; Keiser, M. J.; Shoichet, B. K. Quantifying Biogenic Bias in Screening Libraries. Nat Chem Biol 2009, 5 (7), 479–483.

(17) Irwin, J. J. How Good Is Your Screening Library? Curr Opin Chem Biol 2006, 10 (4), 352–356.

(18) Drewry, D. H.; Macarron, R. Enhancements of Screening Collections to Address Areas of Unmet Medical Need: An Industry Perspective. Curr Opin Chem Biol 2010, 14 (3), 289–298.

(19) Hubbard, R. E. The Role of Fragment-based Discovery in Lead Finding. Fragment-based Drug Discovery Lessons and Outlook 2016, 1–36.

(20) Hall, R. J.; Mortenson, P. N.; Murray, C. W. Efficient Exploration of Chemical Space by Fragment-Based Screening. Prog Biophys Mol Biol 2014, 116 (2–3), 82–91.

(21) Jhoti, H.; Williams, G.; Rees, D. C.; Murray, C. W. The’rule of Three’for Fragment-Based Drug Discovery: Where Are We Now? Nat Rev Drug Discov 2013, 12 (8), 644.

(22) Vukovic, S.; Huggins, D. J. Quantitative Metrics for Drug–Target Ligandability. Drug Discov Today 2018, 23 (6), 1258–1266.

(23) Erlanson, D. A.; McDowell, R. S.; O’Brien, T. Fragment-Based Drug Discovery. J Med Chem 2004, 47 (14), 3463–3482.

(24) Mureddu, L. G.; Vuister, G. W. Fragment-Based Drug Discovery by NMR. Where Are the Successes and Where Can It Be Improved? Front Mol Biosci 2022, 9, 834453.

(25) Davis, B. J.; Giannetti, A. M. The Synthesis of Biophysical Methods In Support of Robust Fragment-Based Lead Discovery. Fragment-based Drug Discovery Lessons and Outlook 2016, 119–138.

(26) Murray, C. W.; Rees, D. C. The Rise of Fragment-Based Drug Discovery. Nat Chem 2009, 1 (3), 187–192.

(27) Shuker, S. B.; Hajduk, P. J.; Meadows, R. P.; Fesik, S. W. Discovering High-Affinity Ligands for Proteins: SAR by NMR. Science (1979) 1996, 274 (5292), 1531–1534.

(28) Labrecque, C. L.; Nolan, A. L.; Develin, A. M.; Castillo, A. J.; Offenbacher, A. R.; Fuglestad, B. Membrane-Mimicking Reverse Micelles for High-Resolution Interfacial Study of Proteins and Membranes. Langmuir 2022, 38 (12), 3676–3686.

(29) Walters, S. H.; Castillo, A. J.; Develin, A. M.; Labrecque, C. L.; Qu, Y.; Fuglestad, B. Investigating Protein-membrane Interactions Using Native Reverse Micelles Constructed from Naturally Sourced Lipids. Protein Science 2023, 32 (11), e4786.

(30) Warschawski, D. E.; Arnold, A. A.; Beaugrand, M.; Gravel, A.; Chartrand, É.; Marcotte, I. Choosing Membrane Mimetics for NMR Structural Studies of Transmembrane Proteins. Biochimica et Biophysica Acta (BBA)-Biomembranes 2011, 1808 (8), 1957–1974.

(31) Dufourc, E. J. Bicelles and Nanodiscs for Biophysical Chemistry. Biochimica et Biophysica Acta (BBA)-Biomembranes 2021, 1863 (1), 183478.

(32) Hagn, F.; Nasr, M. L.; Wagner, G. Assembly of Phospholipid Nanodiscs of Controlled Size for Structural Studies of Membrane Proteins by NMR. Nat Protoc 2018, 13 (1), 79–98.

(33) Klöpfer, K.; Hagn, F. Beyond Detergent Micelles: The Advantages and Applications of Non-Micellar and Lipid-Based Membrane Mimetics for Solution-State NMR. Prog Nucl Magn Reson Spectrosc 2019, 114, 271–283.

(34) Chipot, C.; Dehez, F.; Schnell, J. R.; Zitzmann, N.; Pebay-Peyroula, E.; Catoire, L. J.; Miroux, B.; Kunji, E. R. S.; Veglia, G.; Cross, T. A. Perturbations of Native Membrane Protein Structure in Alkyl Phosphocholine Detergents: A Critical Assessment of NMR and Biophysical Studies. Chem Rev 2018, 118 (7), 3559–3607.

(35) Fuglestad, B.; Kerstetter, N. E.; Bédard, S.; Wand, A. J. Extending the Detection Limit in Fragment Screening of Proteins Using Reverse Micelle Encapsulation. ACS Chem Biol 2019, 14 (10), 2224–2232.

(36) Fuglestad, B.; Kerstetter, N. E.; Wand, A. J. Site-Resolved and Quantitative Characterization of Very Weak Protein–Ligand Interactions. ACS Chem Biol 2019, 14 (7), 1398–1402.

(37) Fuglestad, B.; Marques, B. S.; Jorge, C.; Kerstetter, N. E.; Valentine, K. G.; Wand, A. J. Reverse Micelle Encapsulation of Proteins for NMR Spectroscopy. In Methods in enzymology; Elsevier, 2019; Vol. 615, pp 43–75.

(38) Kielec, J. M.; Valentine, K. G.; Wand, A. J. A Method for Solution NMR Structural Studies of Large Integral Membrane Proteins: Reverse Micelle Encapsulation. Biochimica et Biophysica Acta (BBA)-Biomembranes 2010, 1798 (2), 150–160.

(39) Murray, C. W.; Rees, D. C. Opportunity Knocks: Organic Chemistry for Fragment-based Drug Discovery (FBDD). Angewandte Chemie International Edition 2016, 55 (2), 488–492.

(40) Forcina, G. C.; Dixon, S. J. GPX4 at the Crossroads of Lipid Homeostasis and Ferroptosis. Proteomics 2019, 19 (18), 1800311.

(41) Agmon, E.; Solon, J.; Bassereau, P.; Stockwell, B. R. Modeling the Effects of Lipid Peroxidation during Ferroptosis on Membrane Properties. Sci Rep 2018, 8 (1), 5155.

(42) Ran, Q.; Liang, H.; Gu, M.; Qi, W.; Walter, C. A.; Roberts, L. J.; Herman, B.; Richardson, A.; Van Remmen, H. Transgenic Mice Overexpressing Glutathione Peroxidase 4 Are Protected against Oxidative Stress-Induced Apoptosis. Journal of Biological Chemistry 2004, 279 (53), 55137–55146.

(43) Dixon, S. J.; Lemberg, K. M.; Lamprecht, M. R.; Skouta, R.; Zaitsev, E. M.; Gleason, C. E.; Patel, D. N.; Bauer, A. J.; Cantley, A. M.; Yang, W. S. Ferroptosis: An Iron-Dependent Form of Nonapoptotic Cell Death. Cell 2012, 149 (5), 1060–1072.

(44) Yang, W. S.; SriRamaratnam, R.; Welsch, M. E.; Shimada, K.; Skouta, R.; Viswanathan, V. S.; Cheah, J. H.; Clemons, P. A.; Shamji, A. F.; Clish, C. B. Regulation of Ferroptotic Cancer Cell Death by GPX4. Cell 2014, 156 (1), 317–331.

(45) Ingold, I.; Berndt, C.; Schmitt, S.; Doll, S.; Poschmann, G.; Buday, K.; Roveri, A.; Peng, X.; Freitas, F. P.; Seibt, T. Selenium Utilization by GPX4 Is Required to Prevent Hydroperoxide-Induced Ferroptosis. Cell 2018, 172 (3), 409–422.

(46) Viswanathan, V. S.; Ryan, M. J.; Dhruv, H. D.; Gill, S.; Eichhoff, O. M.; Seashore-Ludlow, B.; Kaffenberger, S. D.; Eaton, J. K.; Shimada, K.; Aguirre, A. J. Dependency of a Therapy-Resistant State of Cancer Cells on a Lipid Peroxidase Pathway. Nature 2017, 547 (7664), 453–457.

(47) Hangauer, M. J.; Viswanathan, V. S.; Ryan, M. J.; Bole, D.; Eaton, J. K.; Matov, A.; Galeas, J.; Dhruv, H. D.; Berens, M. E.; Schreiber, S. L. Drug-Tolerant Persister Cancer Cells Are Vulnerable to GPX4 Inhibition. Nature 2017, 551 (7679), 247–250.

(48) Tsoi, J.; Robert, L.; Paraiso, K.; Galvan, C.; Sheu, K. M.; Lay, J.; Wong, D. J. L.; Atefi, M.; Shirazi, R.; Wang, X. Multi-Stage Differentiation Defines Melanoma Subtypes with Differential Vulnerability to Drug-Induced Iron-Dependent Oxidative Stress. Cancer Cell 2018, 33 (5), 890–904.

(49) Eaton, J. K.; Furst, L.; Cai, L. L.; Viswanathan, V. S.; Schreiber, S. L. Structure–Activity Relationships of GPX4 Inhibitor Warheads. Bioorg Med Chem Lett 2020, 30 (23), 127538.

(50) Bauer, R. A. Covalent Inhibitors in Drug Discovery: From Accidental Discoveries to Avoided Liabilities and Designed Therapies. Drug Discov Today 2015, 20 (9), 1061–1073.

(51) Cozza, G.; Rossetto, M.; Bosello-Travain, V.; Maiorino, M.; Roveri, A.; Toppo, S.; Zaccarin, M.; Zennaro, L.; Ursini, F. Glutathione Peroxidase 4-Catalyzed Reduction of Lipid Hydroperoxides in Membranes: The Polar Head of Membrane Phospholipids Binds the Enzyme and Addresses the Fatty Acid Hydroperoxide Group toward the Redox Center. Free Radic Biol Med 2017, 112, 1–11.

(52) Labrecque, C. L.; Fuglestad, B. Electrostatic Drivers of GPx4 Interactions with Membrane, Lipids, and DNA. Biochemistry 2021, 60 (37), 2761–2772.

(53) Berndt, N.; Yang, H.; Trinczek, B.; Betzi, S.; Zhang, Z.; Wu, B.; Lawrence, N. J.; Pellecchia, M.; Schönbrunn, E.; Cheng, J. Q. The Akt Activation Inhibitor TCN-P Inhibits Akt Phosphorylation by Binding to the PH Domain of Akt and Blocking Its Recruitment to the Plasma Membrane. Cell Death Differ 2010, 17 (11), 1795–1804.

(54) Segers, K.; Sperandio, O.; Sack, M.; Fischer, R.; Miteva, M. A.; Rosing, J.; Nicolaes, G. A. F.; Villoutreix, B. O. Design of Protein–Membrane Interaction Inhibitors by Virtual Ligand Screening, Proof of Concept with the C2 Domain of Factor V. Proceedings of the National Academy of Sciences 2007, 104 (31), 12697–12702.

(55) Caldwell, T. A.; Baoukina, S.; Brock, A. T.; Oliver, R. C.; Root, K. T.; Krueger, J. K.; Glover, K. J.; Tieleman, D. P.; Columbus, L. Low-q Bicelles Are Mixed Micelles. J Phys Chem Lett 2018, 9 (15), 4469–4473.

(56) Baell, J. B.; Holloway, G. A. New Substructure Filters for Removal of Pan Assay Interference Compounds (PAINS) from Screening Libraries and for Their Exclusion in Bioassays. J Med Chem 2010, 53 (7), 2719–2740.

(57) Hajduk, P. J.; Huth, J. R.; Fesik, S. W. Druggability Indices for Protein Targets Derived from NMR-Based Screening Data. J Med Chem 2005, 48 (7), 2518–2525.

(58) Allen, S. J.; Dower, C. M.; Liu, A. X.; Lumb, K. J. Detection of Small-Molecule Aggregation with High-Throughput Microplate Biophysical Methods. Curr Protoc Chem Biol 2020, 12 (1), e78.

(59) Li, C.; Deng, X.; Zhang, W.; Xie, X.; Conrad, M.; Liu, Y.; Angeli, J. P. F.; Lai, L. Novel Allosteric Activators for Ferroptosis Regulator Glutathione Peroxidase 4. J Med Chem 2018, 62 (1), 266–275.

(60) Sedgwick, M.; Cole, R. L.; Rithner, C. D.; Crans, D. C.; Levinger, N. E. Correlating Proton Transfer Dynamics to Probe Location in Confined Environments. J Am Chem Soc 2012, 134 (29), 11904–11907.

(61) Favaro, A.; Sturlese, M. A Novel NMR-Based Protocol to Screen Ultralow Molecular Weight Fragments. J Med Chem 2024.

(62) Yu, H. S.; Modugula, K.; Ichihara, O.; Kramschuster, K.; Keng, S.; Abel, R.; Wang, L. General Theory of Fragment Linking in Molecular Design: Why Fragment Linking Rarely Succeeds and How to Improve Outcomes. J Chem Theory Comput 2020, 17 (1), 450–462.

(63) Van Horn, W. D.; Ogilvie, M. E.; Flynn, P. F. Use of Reverse Micelles in Membrane Protein Structural Biology. J Biomol NMR 2008, 40, 203–211.

(64) Tang, L.; Gamal El-Din, T. M.; Swanson, T. M.; Pryde, D. C.; Scheuer, T.; Zheng, N.; Catterall, W. A. Structural Basis for Inhibition of a Voltage-Gated Ca2+ Channel by Ca2+ Antagonist Drugs. Nature 2016, 537 (7618), 117–121.

(65) Gao, Y.; Cao, E.; Julius, D.; Cheng, Y. TRPV1 Structures in Nanodiscs Reveal Mechanisms of Ligand and Lipid Action. Nature 2016, 534 (7607), 347–351.

(66) Delaglio, F.; Grzesiek, S.; Vuister, G. W.; Zhu, G.; Pfeifer, J.; Bax, A. D. NMRPipe: A Multidimensional Spectral Processing System Based on UNIX Pipes. J Biomol NMR 1995, 6, 277–293.

(67) Lee, W.; Tonelli, M.; Markley, J. L. NMRFAM-SPARKY: Enhanced Software for Biomolecular NMR Spectroscopy. Bioinformatics 2015, 31 (8), 1325–1327.

