## Supplemental Information for "Ligandability at the membrane interface of GPx4 revealed through a reverse micelle fragment screening platform"

**Supplemental Methods**

Micelle, Bicelle, and mmRM Preparation

*N*-dodecyl phosphocholine (DPC), 1,2-dihexanoyl-sn-glycero-3-phosphocholine (DHPC) and 1,2-dimyristoyl-sn-glycero-3-phosphocholine (DMPC), and 1,2-dilinoleoyl-*sn*-glycero-3-phosphocholine (DLPC), all purchased from Avanti Polar Lipids (Birmingham, AL) were vacuum dried overnight to remove residual water prior to membrane model construction. DPC micelles were formed aby weighing 25 mM total surfactant into a vial and resuspending in GPx4 NMR buffer (20 mM Bis-Tris pH 6.0, 100 mM NaCl, and 10 mM DTT). Micelles spontaneously form with vortexing and no other steps are required. DHPC:DMPC bicelles are created at a 1:1 ratio of 25 mM surfactants. Briefly, DMPC is weighed into a glass vial and then dissolved in chloroform. The chloroform is dried off under a stream of nitrogen while rotating the vial to ensure a thin film is formed. The DHPC fraction was weighed and then dissolved in GPx4 buffer before adding the solution to the film of DMPC. The slurry underwent 5 freeze-thaw cycles, alternating between an ice bucket and a 70 °C water bath, for 5 minutes with vigorous vortexing in between. After 5 cycles, the bicelle solution was visually clear. DLPC:DPC mmRMs were formed by weighing the surfactant to a concentration of 75 mM and a ratio of 1:1. Pentane was added along with 200 mM 1-hexanol (Sigma-Aldrich, St. Louis, MO) to form a slurry. GPx4 NMR buffer was added for a water loading ($W_0$) of 20 defined as:

$$W_0 = \frac{[water]}{[surfactant]}$$

Upon buffer addition, hexanol was titrated 200 mM at a time until 1 M at which point visual sample clarity was reached.

Dynamic Light Scattering (DLS)

All membrane models, DPC micelles, DHPC:DMPC bicelles, and DLPC:DPC mmRMs were evaluated by DLS to determine their validity as a membrane model for the fragment screen. For DLS measurements, the mmRM samples were formed as before, but hexane (Thermo Fisher Scientific, Waltham, MA) replacing pentane to avoid rapid evaporation. Experiments were performed using a Malvern Zetasizer Nano-S instrument. Published viscosity and dielectric constant parameters were used for binary mixtures of hexane and hexanol for the mmRM samples[1,2] and values for water were used for bicelle and micelle samples[3,4]. The samples were loaded into a quartz cuvette and held at 25 °C throughout the course of the measurement. Each sample was scanned 12-15 times to complete one measurement, and each was completed in triplicate to generate an average size of the population with error. After the micelle and bicelle measurements were collected, 5% DMSO was added to the solution and vortexed before recollection. For the fragment mixture experiments with micelles and mmRMs, 10 mixtures each composed of 10 fragments were selected and dried to remove the DMSO storage buffer. The mixtures were incorporated into 10 separate micelle and mmRM samples and mixed for an hour before measurement. All samples were visually assessed to evaluate fragment solubility before measurement.

Protein Expression and Purification

Truncated U46G-GPx4 with a TEV cleavable poly-histidine tag was inserted into vector pNIC28-Bsa4 and was a gift from Nicola-Burgess-Brown (Addgene plasmid #38797). The plasmid was transformed into BL21 (DE3) E. coli and grown on LB-agar plates containing kanamycin. Glycerol stocks were made from the colonies and used to inoculate overnight cultures of M9 minimal media grown at 37 °C. The following morning, the overnight growths were pelleted by centrifugation and used to seed 1L growths of M9 media with $^{15}$N-ammonium chloride was added to isotopically label the proteins of interest. The growths were grown to an $OD_{600}$ of 0.800 before induction with 1 mM isopropyl-β-D-1-thiogalactopyranoside (IPTG) and grown for roughly 18 hours at 30 °C. After 18 hours, the cells were harvested by centrifugation for lysis and purification.

GPx4 was lysed and prepared for NMR analysis as previously described.[5] Briefly, $^{15}$N-labeled protein was lysed by sonication and purified using a Ni-NTA resin column. Five column volumes of wash buffer (100 mM Tris pH 7.4, 300 mM NaCl, 50 mM imidazole, and 5 mM DTT) were flowed over the resin to remove non-specific binders to the nickel beads. The wash step was followed by three column volumes of elution buffer (all buffer components were the same but imidazole was increased from 50 mM to 300 mM). The cleavable His-tag was removed with TEV protease and then repurified for final experimentation. The NMR buffer for GPx4 was 20 mM Bis-Tris pH 6.0, 100 mM NaCl, and 20 mM DTT.

### Supplemental Figures

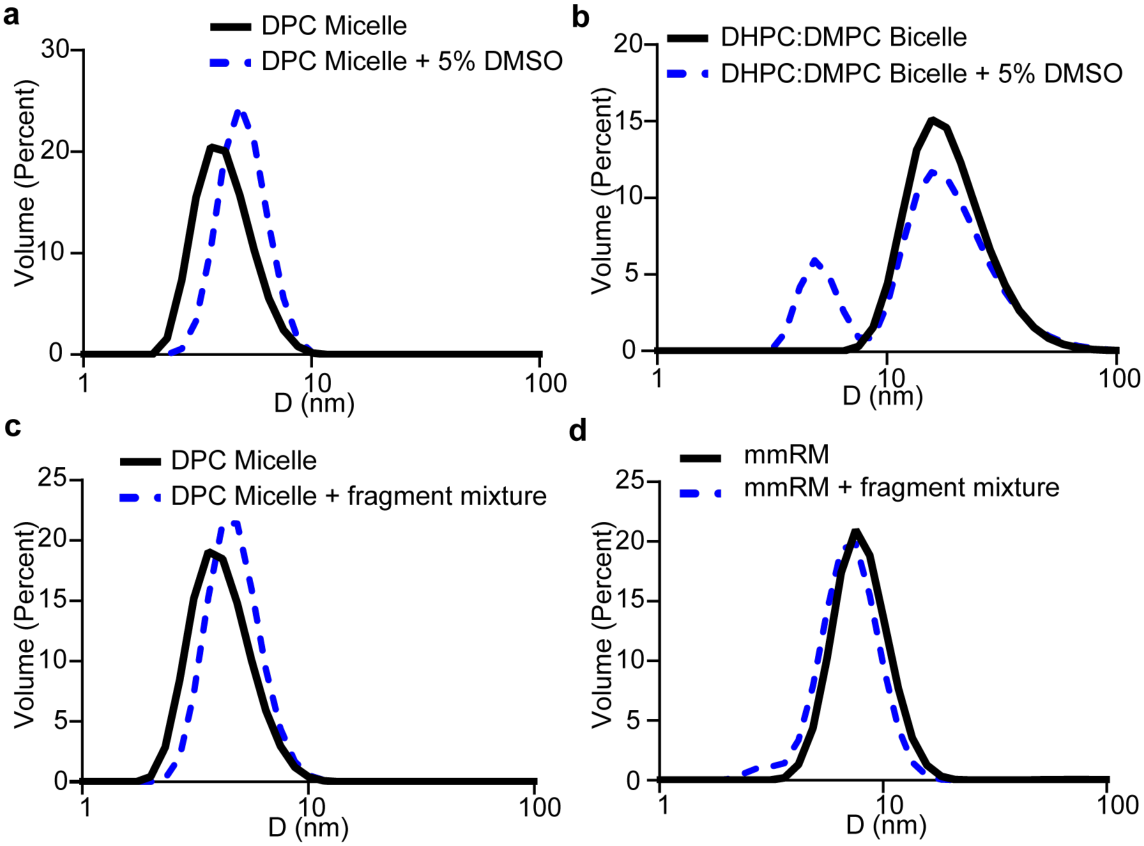


**Figure S1.** Compatibility of membrane models for fragment screening assessed by DLS measurements. Addition of DMSO to micelle and bicelle models to allow for fragment solubilization in the bulk aqueous phase causes model perturbations. **a)** 25mM DPC micelle before and after the addition of 5% DMSO. Micelle size increased from 4.27 ± 0.040 nm to 5.11 ± 0.030 nm, indicating incorporation of DMSO. **b)** 1:1 (25mM) DHPC:DMPC bicelle before and after the addition of 5% DMSO. Bicelle size increased from 20.06 ± 1.54 nm to 21.33 ± 0.50 nm with DMSO as well as formation of a second population at 5.21 ± 1.05 nm. **c)** 25 mM DPC micelle before and after the addition of pre-dried fragment mixture. Prior to the fragment addition, the micelle size was 4.29 ± 0.13 nm and increased to 4.83 ± 0.42 nm with the addition of the fragment mixture. **d)** The addition of a fragment mixture to a 1:1 (75mM) DLPC-DPC mmRM caused a decrease in the average size of the mmRM from 8.23 ± 0.64 nm to 7.49 ± 0.88 nm with the addition of fragment mixtures. For both **c)** and **d)** measurements with fragment, the same 10 mixtures were used to collect an average size perturbation.

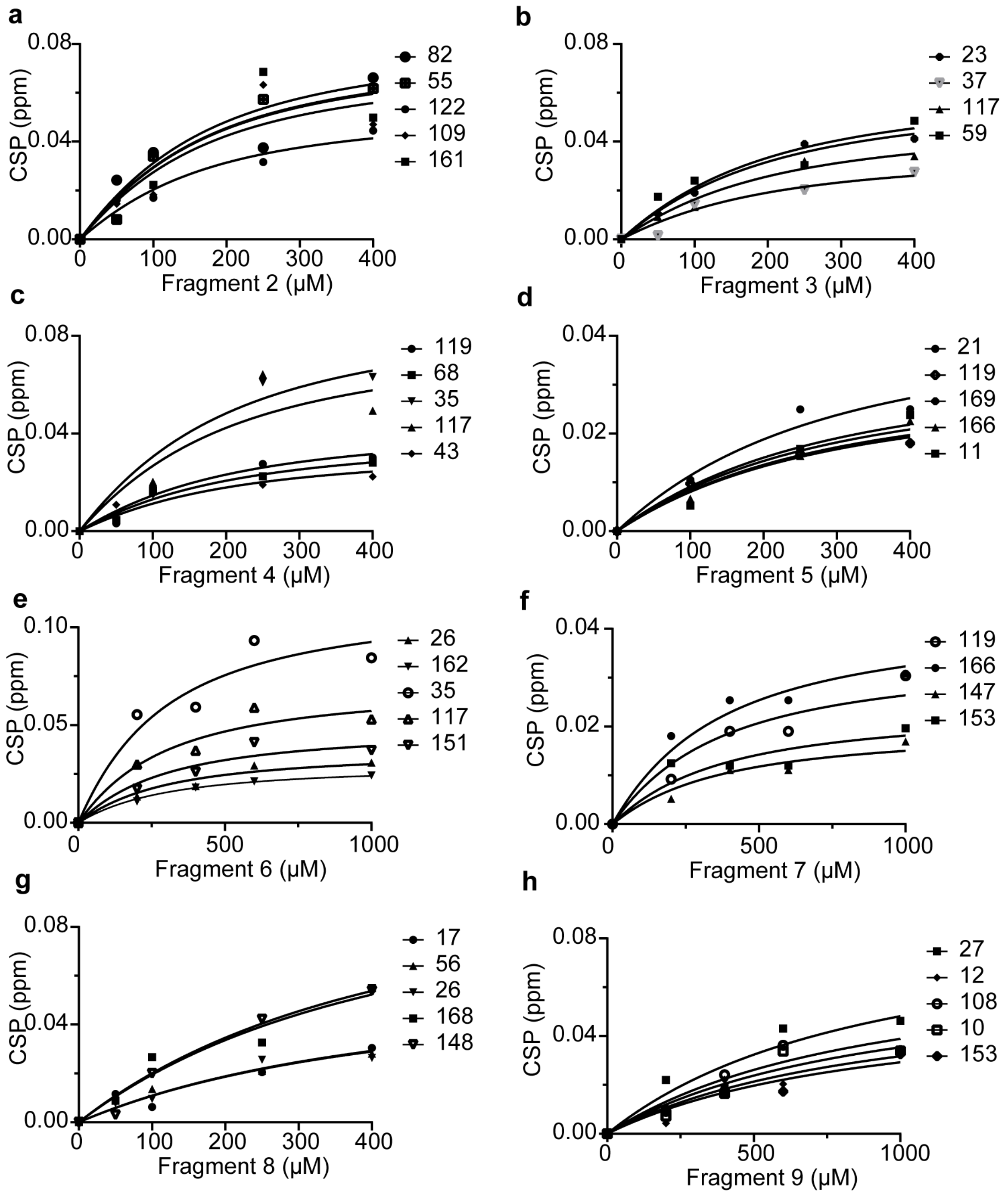


**Figure S2:** All titrations for fragment hit characterization. All curves shown were used for apparent $K_d$ analysis. The top concentration used for the titrations was either 400 µM (a, b, c, d, g) or 1 mM (e, f, h). Resonances used for analysis were selected by identifying the top-sifting resonances at the highest fragment concentration. These were resonances with a CSP 1σ above the average CSP for all observable resonances. The top-shifters were then fit to individual $K_d$ curves and anything with a $R^2 < 0.85$ was removed. The remaining resonances were then included in a global fit to determine the overall apparent $K_d$ for the fragment titration. All apparent $K_d$s are listed in **Table 1**.

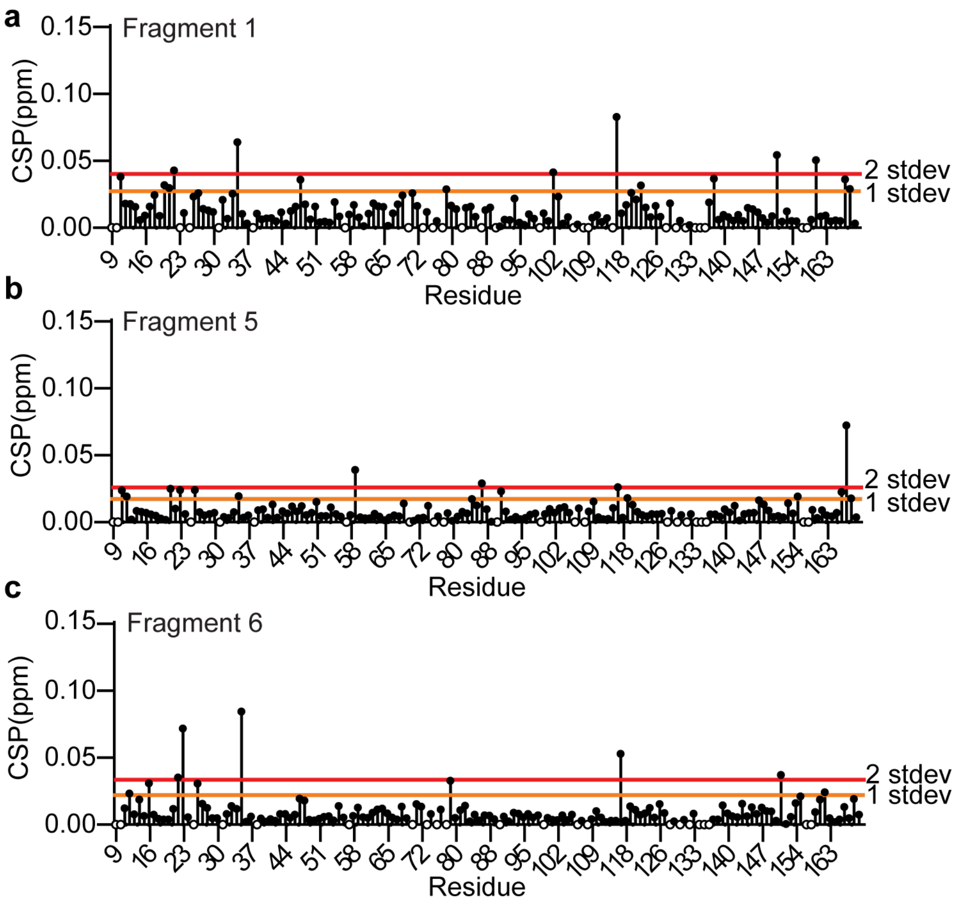


**Figure S3.** CSP analysis of GPx4 embedded in mmRMs bound to fragments. 400 µM **a)** fragment 1, **b)** fragment 5, and **c)** fragment 6 were used. CSPs were calculated between [$^{1}$H-$^{15}$N] HSQCs of DLPC:DPC encapsulated GPx4 in the presence and absence of fragment. Each mmRM was constructed of 75 mM DLPC:DPC at a 1:1 ratio with 1 M hexanol, $W_0 = 20$, and dissolved in pentane. The buffer used in the $W_0$ was 100 mM Bis-Tris pH 6.0, 100 mM NaCl, and 10 mM DTT. The average CSP for the observable resonances was calculated and used to determine CSP cutoffs 1 (orange line) and 2 (red line) σ above the average. Resonances above the 1 σ were used to calculate apparent $K_d$s and to map onto the crystal structure of GPx4.

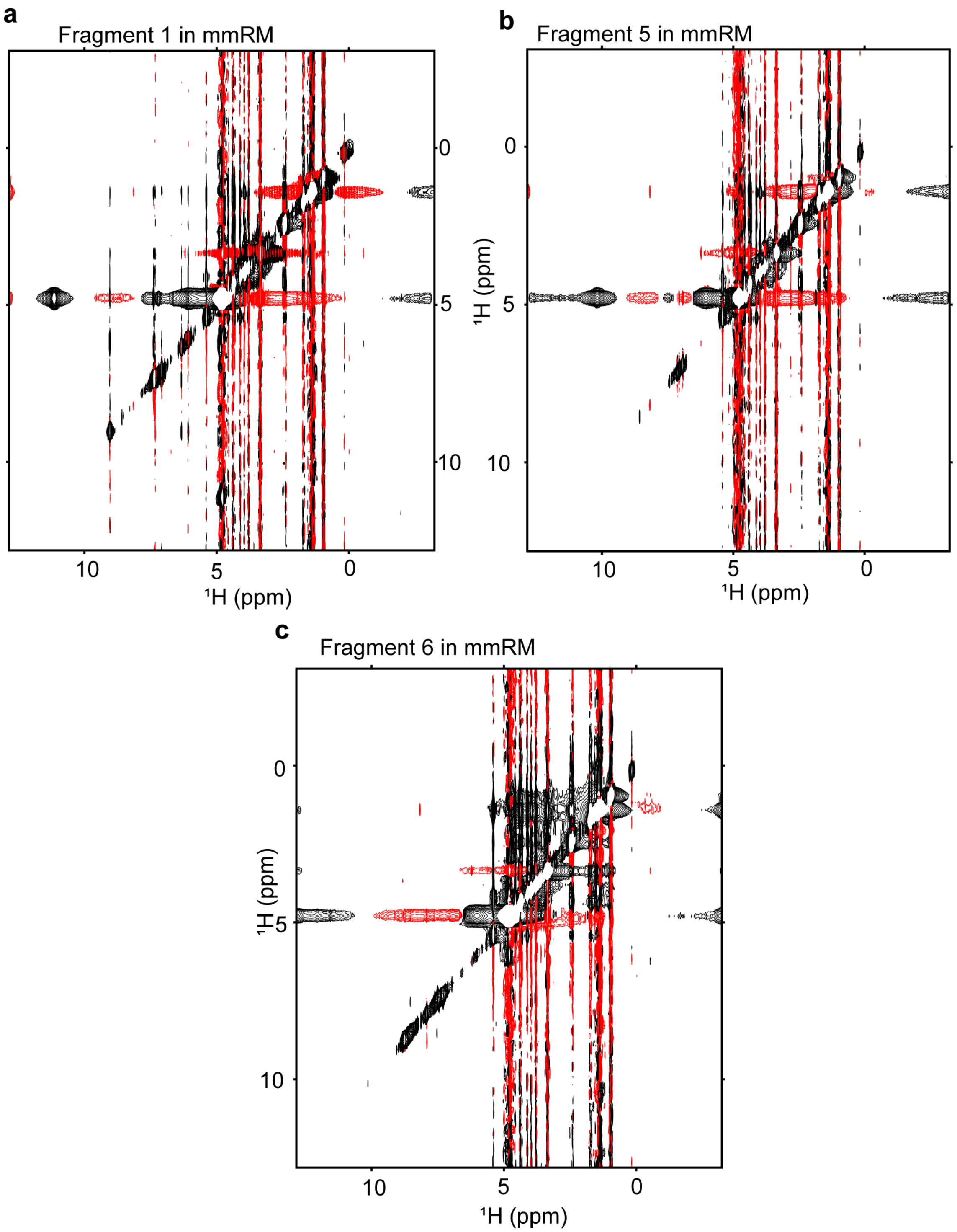


**Figure S4.** Full [$^1$H-$^1$H] NOESY spectra of 5 mM **a)** fragment 1, **b)** fragment 5, and **c)** fragment 6 depicting the interaction between the fragments and components of the DLPC:DPC mmRMs. mmRM were constructed with 75 mM DLPC:DPC and a molar ratio of 1:1, 1 M d-hexanol, and d-pentane to reduce the solvent peaks. The $W_0$ was 20 and composed of GPx4 NMR buffer. Negative contours are shown in red and positive contours are shown in black. Contour levels were raised to near the noise level surrounding NOE crosspeaks, accounting for the enhanced artifacts.

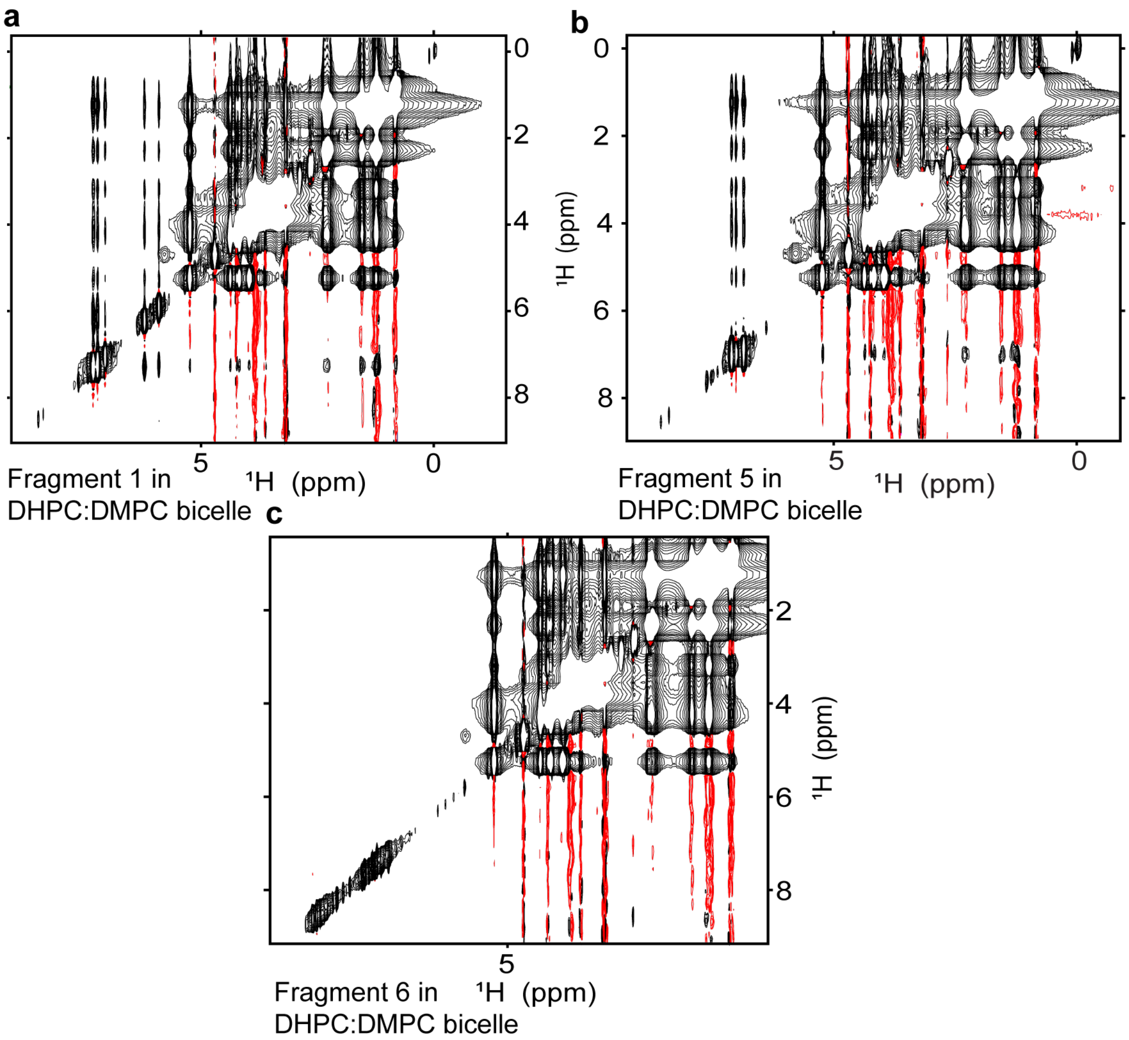


**Figure S5.** Full [1H-1H] NOESY spectra of 5 mM **a)** fragment 1, **b)** fragment 5, and **c)** fragment 6 showing the interaction between the fragments and DHPC:DMPC bicelles. Bicelles were constructed of 1:1 (25 mM each) DHPC:DMPC and dissolved in NMR buffer made in 100% $D_2O$ to reduce the water peak. Negative contours are shown in red and positive contours are shown in black. Contour levels were raised to near the noise level surrounding the NOE crosspeaks, accounting for the enhanced artifacts.

| Analog Number | Analog Structure | Apparent $K_d$ (μM) | Predicted cLogD (pH 6.0) |
|---|---|---|---|
| 1 | 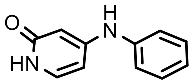 | 105 ± 30 | 1.05 |
| 1.1 | 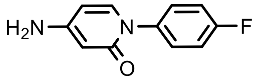 | > 400 | 1.01 |
| 1.2 | 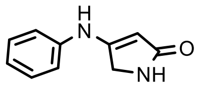 | > 400 | 0.34 |
| 1.3 | 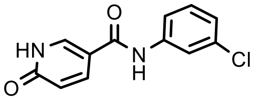 | > 400 | 1.49 |
| 1.4 | 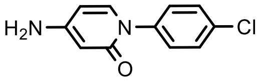 | > 400 | 1.47 |
| 1.5 | 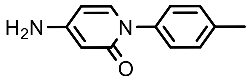 | > 400 | 1.38 |

**Table S1.** Apparent $K_d$s and predicted cLogDs of titrated analogs of fragment 1.

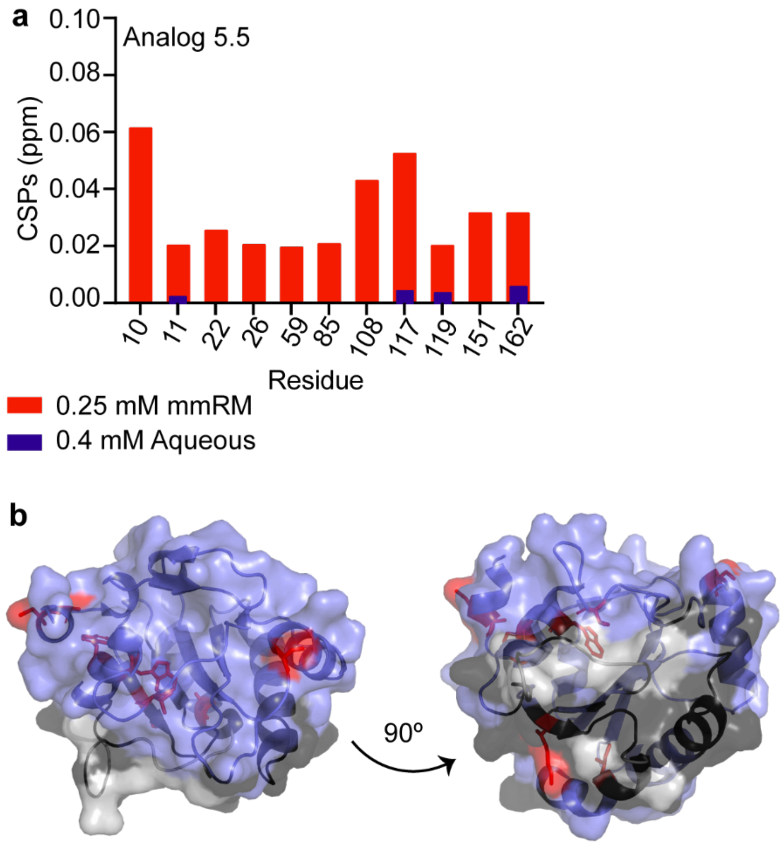


**Figure S6.** CSPs of mmRM embedded GPx4 upon addition of Analog 5.5. a) Top-shifting resonances from [$^1$H-$^{15}$N] HSQC experiments were identified by calculating the CSP between GPx4 encapsulated in mmRM and GPx4 in mmRMs with 400 μM analog 5.5. 1 σ above of the average were isolated as the top-shifters (red bars). These shifting resonances were then compared to the equivalent aqueous experiment which consisted of apo GPx4 with 1.5% DMSO and GPx4 with 400 μM analog 5.5 an 1.5% DMSO (blue bars). The increased DMSO was to ensure the analog was completely solubilized prior to beginning the experiment. **b)** The top-shifting resonances were then mapped on a crystal structure of GPx4 with the membrane interface in DPC micelles shown in black, residues with little or no shifting are shown in blue, missing resonances in grey, and analog interacting residues shown with red sticks.

### Supplemental References


(1) Franjo, C.; Jimenez, E.; Iglesias, T. P.; Legido, J. L.; Paz Andrade, M. I. Viscosities and Densities of Hexane+ Butan-1-Ol,+ Hexan-1-Ol, And+ Octan-1-Ol at 298.15 K. *J Chem Eng Data* **1995**, *40* (1), 68–70.

(2) Singh, R. P.; Sinha, C. P. Dielectric Behavior of the Binary Mixtures of N-Hexane with Toluene, Chlorobenzene and 1-Hexanol. *J Chem Eng Data* **1982**, *27* (3), 283–287.

(3) Bonincontro, A.; Cametti, C.; Sesta, B. Density, Viscosity and Dielectric Constant of Aqueous Solutions of Triglycine and Tetraglycine. *Zeitschrift für Naturforschung A* **1978**, *33* (4), 462–467.

(4) Dembek, M.; Bocian, S. Pure Water as a Mobile Phase in Liquid Chromatography Techniques. *TrAC Trends in Analytical Chemistry* **2020**, *123*, 115793.

(5) Labrecque, C. L.; Fuglestad, B. Electrostatic Drivers of GPx4 Interactions with Membrane, Lipids, and DNA. *Biochemistry* **2021**, *60* (37), 2761–2772.
